# DOT1L-AF10–mediated H3K79me3 promotes NF-κB p65–dependent inflammatory activation in endothelial cells

**DOI:** 10.64898/2026.03.20.713137

**Authors:** Yash T Katakia, Ritobrata Bhattacharyya, Sushmitha Duddu, Nivedha Suresh, Srinjoy Chakraborty, Nehal Gupta, Suchit Chebolu, Praphulla Chandra Shukla, Syamantak Majumder

## Abstract

DOT1L-catalyzed H3K79 methylation is a hallmark of actively transcribed genes and has been extensively studied in developmental and disease contexts. While DOT1L inhibition has emerged as a promising therapeutic strategy in cancer, its role in pro-atherogenic endothelial inflammation remains unclear. To investigate this, we utilized an *in vivo* partial carotid artery ligation model and observed increased DOT1L expression and H3K79me3 level. Consistently, *in vitro* studies employing a 3D-printed human coronary artery model and TNF-α stimulation corroborated these results, showing elevated DOT1L expression and H3K79me3 deposition, while levels of H3K79me and me2 remained unchanged. Further analyses identified key DOT1L-containing complex (DotCom) components, AF10 and AF9 (upregulated) and AF17 (downregulated), as contributors to the enhanced H3K79me3 landscape. CUT&RUN sequencing showed prominent H3K79me3 enrichment at the *RELA* (NF-κB p65) promoter, corresponding with increased NF-κB p65 expression and activation. Notably, inhibition/knockdown of the methyltransferase DOT1L or overexpression of the demethylase FBXL10 significantly reduced H3K79me3 levels, thereby suppressing NF-κB p65 expression and attenuating endothelial inflammation, independent of canonical NF-κB p65 activation. These findings establish DOT1L-mediated H3K79me3 as a crucial epigenetic regulator of endothelial inflammation, highlighting a potential therapeutic avenue for mitigating NF-κB p65-driven pro-atherogenic endothelial dysfunction.

## Introduction

Histone modifications, particularly methylation, are essential epigenetic regulators of gene expression, shaping diverse cellular processes. Among these, DOT1L (disruptor of telomeric silencing 1-like) has garnered considerable attention owing to its catalytic activity that facilitates the methylation of histone H3 lysine 79 (H3K79), a mark predominantly associated with transcriptionally active chromatin (1, 2). While DOT1L-mediated H3K79 methylation is well-characterized in leukemogenesis (3, 4), emerging research has begun to highlight its broader regulatory roles in developmental contexts (5–7) and disease settings (8, 9), particularly in vascular biology. Notably, DOT1L has been shown to regulate lymphatic endothelial progenitors, playing a critical role in normal lymphatic development and function (10). Moreover, studies linking DOT1L-catalyzed H3K79 methylation to inflammatory diseases (11, 12), suggest that DOT1L represents a promising therapeutic target in vascular pathologies.

Atherosclerosis is a progressive chronic inflammatory disease marked by lipid accumulation and plaque formation within arterial walls, primarily initiated by endothelial dysfunction and sustained by persistent pro-inflammatory signaling (13). The transcription factor NF-κB p65 is central to the regulation of inflammatory gene expression in endothelial cells (EC), becoming activated in response to stimuli such as TNF-α. Upon activation, NF-κB p65 induces the expression of adhesion molecules, cytokines, and other pro-inflammatory mediators that promote vascular inflammation and plaque development (14). Although the canonical NF-κB signaling pathway has been extensively studied, the influence of epigenetic modifications–specifically H3K79 methylation–on NF-κB p65 expression and its atherogenic activity in EC remains less understood.

In this study, we demonstrate that EC exposed to disturbed flow (D-Flow) and TNF-α exhibit upregulated DOT1L expression and increased H3K79me3 enrichment at the promoter region of the *RELA* (NF-κB p65) gene. This epigenetic modification is linked to enhanced NF-κB p65 expression, thereby sensitizing the endothelium to an inflammatory response. Importantly, the inhibition of DOT1L-catalyzed H3K79me3 resulted in decreased NF-κB p65 protein levels and attenuated endothelial inflammation. These findings suggest that DOT1L-mediated H3K79me3 serves as a key epigenetic regulator of NF-κB p65 expression, acting independently of its canonical activation pathway. DOT1L and H3K79me3 play a pivotal role in governing EC responses to pro-atherogenic signals by modulating the transcriptional landscape of inflammatory genes.

Given the fundamental role of endothelial dysfunction in the onset and progression of atherosclerosis, targeting DOT1L and its downstream epigenetic markers, such as H3K79me3, presents a promising therapeutic strategy to mitigate NF-κB p65-driven vascular inflammation. These insights reveal the potential of DOT1L inhibitors in controlling endothelial inflammation and halting the pathophysiological processes that drive atherosclerosis.

## Results

### *In vivo* exposure to disturbed flow increases endothelial DOT1L and H3K79me3 levels

DOT1L-catalyzed H3K79 methylations are well known for decorating actively transcribed genes. The involvement of DOT1L in various physiological and pathological processes, including angiogenesis, cardiomyocyte differentiation, neural development and neurological disorders, the transition of vascular smooth muscle cells (VSMC) phenotype, and cancers has been extensively studied (3, 6, 7, 10, 12). However, the specific role of DOT1L and H3K79 methylation in atherogenic endothelial dysfunction remains elusive. Thus, we aimed to investigate the effects of reduced WSS or D-Flow on the endothelial expressions of DOT1L and H3K79 methylation. To address this, we employed a partial carotid artery ligation model in HFD-fed C57BL/6 mice (15). Four weeks post-surgery, the mice were sacrificed, and carotid artery tissues, both ligated as well as contralateral unligated (control), were isolated and processed for immunohistochemical analysis. Hematoxylin and eosin staining of the histological sections revealed neointima formation in the ligated carotid artery, while no such changes were observed in the unligated control artery (Supplementary Figure 1A). Additionally, immunofluorescence staining of the histological sections revealed elevated endothelial expressions of DOT1L (Figure 1A) and its methylation product, H3K79me3 (Figure 1B), in the ligated arteries. A recent study indicated that VSMC within neointima or atherosclerotic plaques exhibit upregulated DOT1L expression (12). Consistent with these findings, our evaluation revealed a marked increase in DOT1L expression in the VSMC of the neointimal region of the ligated artery compared to the unligated control (Supplementary Figure 1B). To further evaluate the spatial distribution of DOT1L and its associated histone mark H3K79me3 under distinct hemodynamic conditions, rat aortas were subjected to *en face* immunostaining to assess their expression in regions exposed to disturbed flow (D-flow; lesser curvature) and stable laminar flow (S-flow; greater curvature). EC located in D-flow regions exhibited markedly elevated DOT1L expression (Figure 1C) along with a corresponding increase in H3K79me3 levels (Figure 1D) compared to EC in S-flow regions. In addition, en face staining was performed on intact aortic rings following ex vivo treatment with TNF-α (24 h). TNF-α exposure resulted in a significant upregulation of H3K79me3 within the endothelial monolayer (Figure 1E), further supporting the association between inflammatory stimulation and enhanced DOT1L-mediated histone methylation.

**Figure 1.**
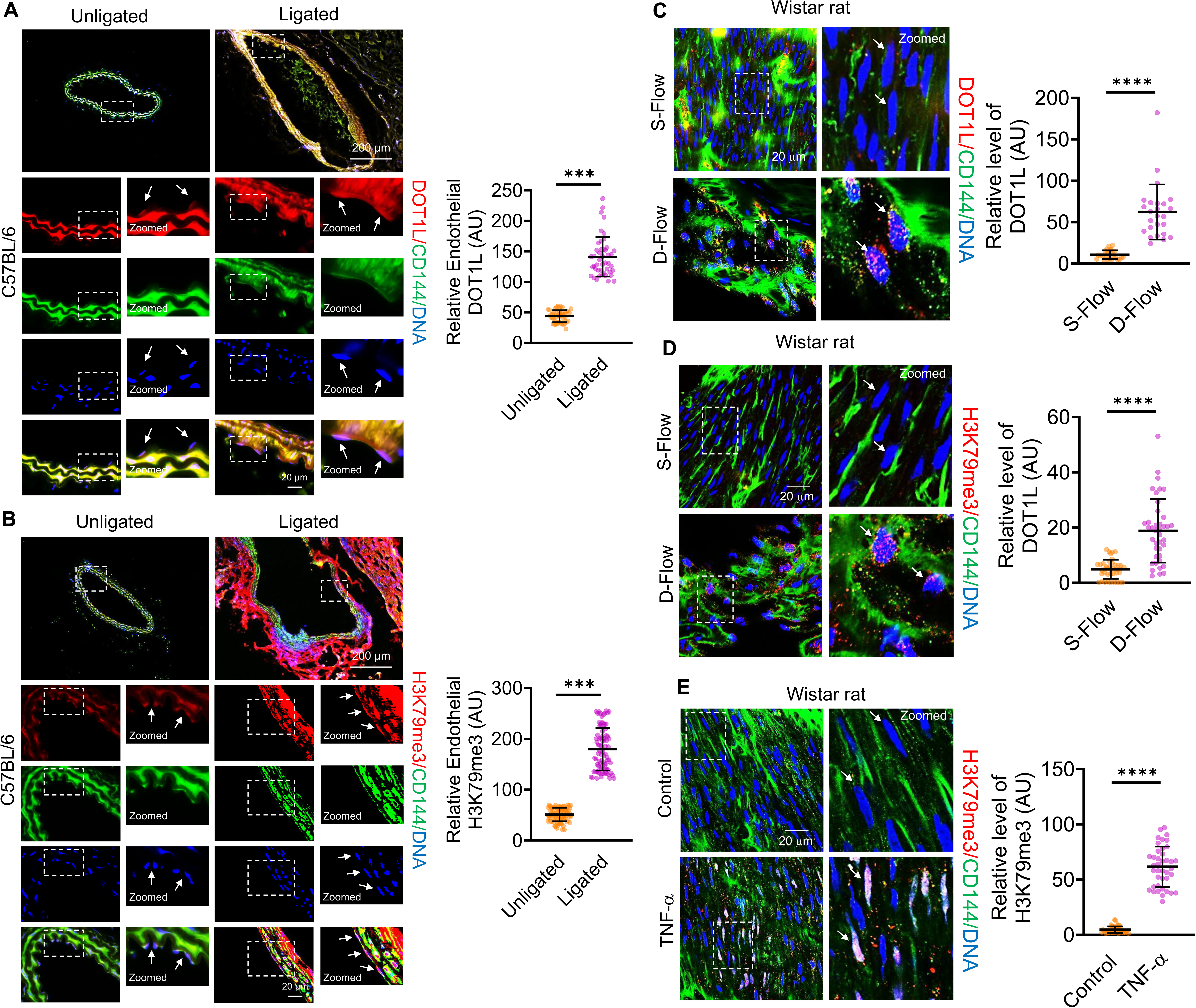
D-Flow induces endothelial DOT1L expression and H3K79me3 enrichment *in vivo*. (A-B) Immunofluorescence staining of histological sections from partially ligated (D-Flow) and contralateral unligated (S-Flow control) carotid arteries of HFD-fed C57BL/6 mice (n = 3), performed four weeks post-surgery, showing endothelial expression of DOT1L (A) and H3K79me3 (B). EC are marked by CD144 (green), nuclei by DAPI (blue), and target signals (DOT1L or H3K79me3) in red. Fluorescence intensities (AU) in individual EC (dots) from three biological replicates are indicated together with the mean. A total of ≥ 45 EC were analyzed per condition. (C-E) *En face* preparations were double stained with anti–VE-cadherin (green, EC marker) and an anti-DOT1L antibody (red, C) or H3K79me3 antibody (red, D, E) in both lesser (disturbed flow [D-flow]) and greater (steady laminar flow [S-flow]) curvature areas including TNF-α treated aortic tissue (E). Images were obtained from the luminal surface of the aorta. The levels of nuclear EZH2 and H3K27me3 were analyzed using Image J software. (n = 3) Magnification: x10, x63; Scale: 200 µm, 20 µm. Data are presented as mean ± SD. *p < 0.05, **p < 0.01, ***p < 0.001, ****p < 0.0001 by Welch’s t-test.

### Elevated DotCom associated gene expressions enhance H3K79me3 levels in endothelial cells exposed to either disturbed flow or TNF-α *in vitro*

After evaluating DOT1L and its product H3K79me3 in an *in vivo* D-Flow model, we sought to corroborate these findings in an *in vitro* setting. To assess changes in DotCom complex transcript levels under inflammatory conditions, we analyzed publicly available RNA-seq data from TNF-α–treated HUVECs (0, 4, and 24 h; GSE121958). Several DotCom-associated genes, including *DOT1L*, *AF9*, *AF10*, *AF17*, and *ENL*, exhibited increased expression, with the most pronounced upregulation observed at 4 h following TNF-α stimulation (Figure 2A-B). Notably, transcript levels of *AF9*, *AF10*, and *ENL* showed a modest decline after 24 h of TNF-α exposure, suggesting a dynamic and time-dependent transcriptional regulation of DotCom components in response to inflammatory signaling (Figure 2A-B). To this end, we utilized 3D-printed human coronary artery microchannels to subject EC to *in vivo*-like D-Flow conditions (16) and TNF-α stimulation to induce an inflammatory endothelial phenotype (17). Under these conditions, we observed an elevation in H3K79me3 enrichment levels in EA.hy926 cells exposed to D-Flow (Figure 2C, Figure 2D, Figure 2E) and TNF-α (Supplementary Figure 2A), as well as in HUVEC treated with TNF-α (Supplementary Figure 2B). Interestingly, TNF-α treatment did not alter the level of H3K79me (Supplementary Figure 2C) and H3K79me2 (Supplementary Figure 2D) in cultured EC. Furthermore, our analysis revealed a significant upregulation of DOT1L expression in EC subjected to either D-Flow (Figure 2F) or TNF-α (Figure 2G, Supplementary Figure 2E).

**Figure 2.**
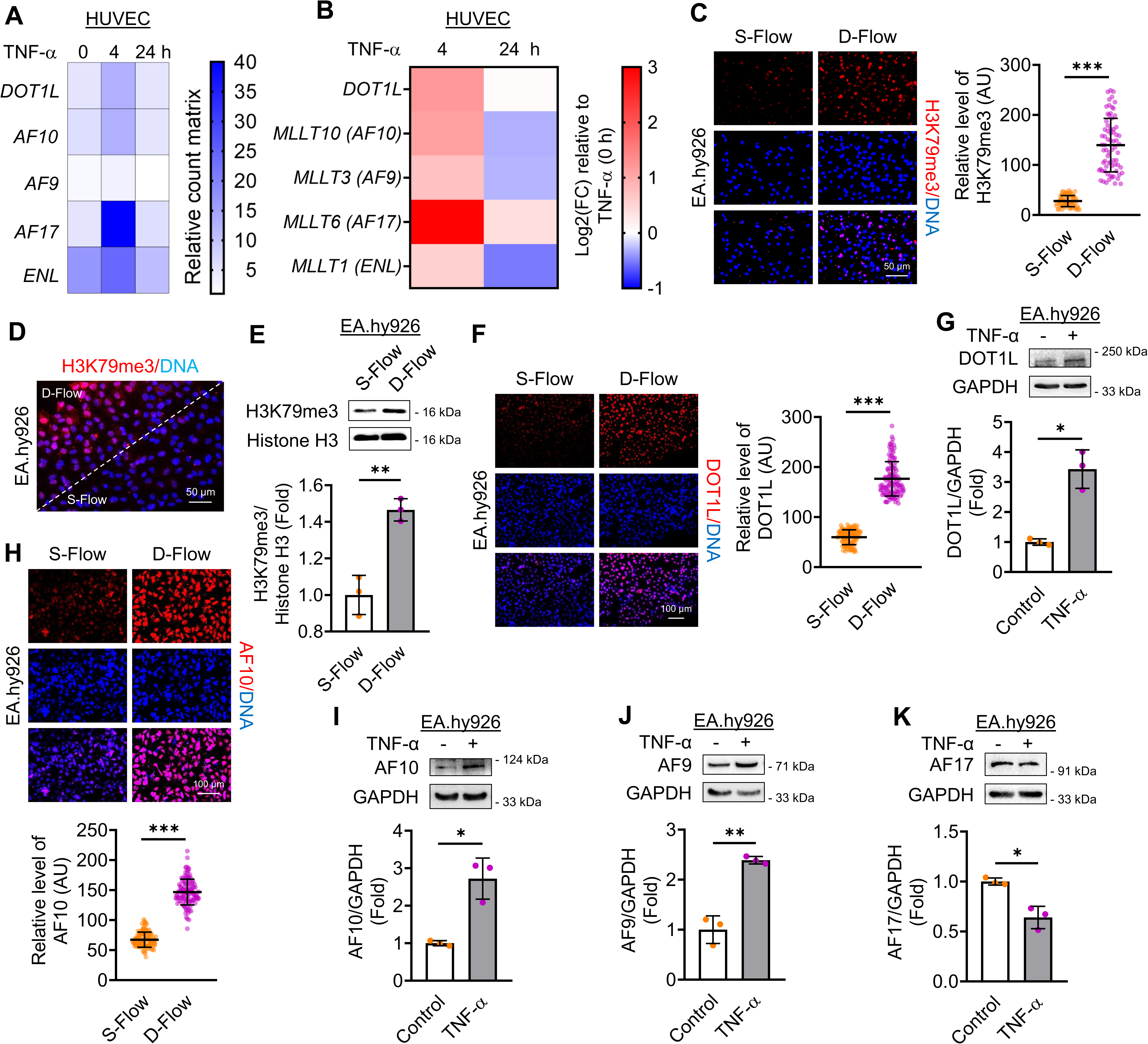
D-Flow and TNF-α modulate DotCom protein expression to promote endothelial H3K79me3 *in vitro*. (A-B) Heatmap derived from RNA-seq data (GEO: GSE121958) showing relative count matrix (E) and log₂ fold change (F) in DOT1L complex-associated genes following TNF-α stimulation (4 and 24 h) to HUVEC. (C-D) Immunofluorescence staining of EA.hy926 exposed to D-Flow (4 h; n = 3), showing H3K79me3 enrichment. Nuclei are stained with DAPI (blue), and H3K79me3 is shown in red. Fluorescence intensities (AU) in individual EC (dots) from three independent experiments are indicated together with the mean. A total of ≥ 60 EC were analyzed per condition. Magnification: x40; Scale: 50 µm. (E) Immunoblot analysis of H3K79me3 levels in EA.hy926 exposed to S- and D-Flow (4 h; n = 3). (F) Immunofluorescence staining of EA.hy926 exposed to D-Flow (4 h; n = 4), showing DOT1L expression (red). Nuclei are stained with DAPI (blue). Fluorescence intensities (AU) in individual EC (dots) from four independent experiments are indicated together with the mean. A total of ≥ 115 EC were analyzed per condition. Magnification: x20; Scale: 100 µm. (G) Immunoblot analysis of DOT1L expression in EA.hy926 treated with TNF-α (10 ng/ml, 24 h; n = 3). (H) Immunofluorescence staining of EA.hy926 exposed to D-Flow (4 h; n = 3), showing AF10 expression in red and nuclei in blue. Fluorescence intensities (AU) in individual EC (dots) from three independent experiments are indicated together with the mean. A total of ≥ 120 EC were analyzed per condition. Magnification: x20; Scale: 100 µm. (I-K) Immunoblot analyses of AF10 (I), AF9 (J), and AF17 (K) in EA.hy926 treated with TNF-α (10 ng/ml, 24 h; n = 3). Data are presented as mean ± SD. *p < 0.05, **p < 0.01, ***p < 0.001 by Welch’s t-test.

Having established that both D-Flow and TNF-α exposures promote DOT1L-catalyzed H3K79me3 in EC, we next sought to investigate the expression of other DOT1L-containing Complex (DotCom) proteins. Specifically, we assessed the expression of DOT1L-binding partners, including AF10/MLLT10, AF9/MLLT3, and AF17/MLLT6. AF10, a well-characterized DOT1L co-factor (18), recruits the DotCom by recognizing unmodified H3K27(19). Our results demonstrated a significant increase in AF10 expression in EC exposed to D-Flow (Figure 2H) or TNF-α (Figure 2I, Supplementary Figure 2F). Additionally, AF9, another DOT1L-binding partner, is known to bind acetylated histone H3 and facilitate DOT1L recruitment (20). EC treated with TNF-α exhibited enhanced AF9 expression (Figure 2J, Supplementary Figure 2F). In contrast, AF17, a DotCom protein involved in promoting DOT1L nuclear export and negatively regulating H3K79 methylation (21, 22), showed a marked reduction in expression under both D-Flow and TNF-α stimulation (Figure 2K, Supplementary Figure 2G-H).

### TNF-α treatment caused enhanced H3K79me3 deposition at the *RELA* / *NF-κB p65* gene promoter in endothelial cells

To characterize the genome-wide distribution of H3K79me3 during endothelial inflammation, we performed CUT&RUN sequencing in HUVECs treated with or without TNF-α. Chromosome-wide coverage analysis showed broadly conserved H3K79me3 enrichment across all chromosomes, with TNF-α stimulation producing focal changes in signal intensity without large-scale chromosomal redistribution (Supplementary Figure 3A). Genome-wide feature annotation revealed that H3K79me3 peaks were predominantly localized to distal intergenic regions in both conditions, followed by intronic and promoter-associated regions, and overall genomic distributions remained similar between untreated and TNF-α-treated cells. TNF-α exposure resulted in modest increases in peaks associated with the 5′ and 3′ untranslated regions (UTRs) and subtle enrichment within promoter-proximal bins, indicating selective changes in regulatory element occupancy (Figure 3A). Metagene analysis centered on transcription start sites (TSS) demonstrated a pronounced promoter-associated enrichment of H3K79me3 in both conditions, with TNF-α treatment modestly increasing signal intensity around the TSS without altering the overall distribution pattern, as supported by aggregate profiles and heatmap visualization (Supplementary Figure 3B). Consistent with these observations, distance-to-TSS analysis showed that the majority of H3K79me3-associated loci were positioned at distal regions, predominantly within the 10–100 kb interval from the TSS, whereas TNF-α stimulation modestly increased the proportion of peaks within proximal bins (0–10 kb) without altering the overall positional distribution (Figure 3B). UpSet plot analysis showed greater diversity and complexity of genomic feature intersections in TNF-α-treated cells, including increased co-occurrence of H3K79me3 peaks at the promoter, intronic, and upstream regions (Figures 3C-D; Supplementary Figure 3B-C). This suggests a more intricate regulatory landscape under inflammatory conditions, with possible long-range enhancer-promoter interactions.

**Figure 3.**
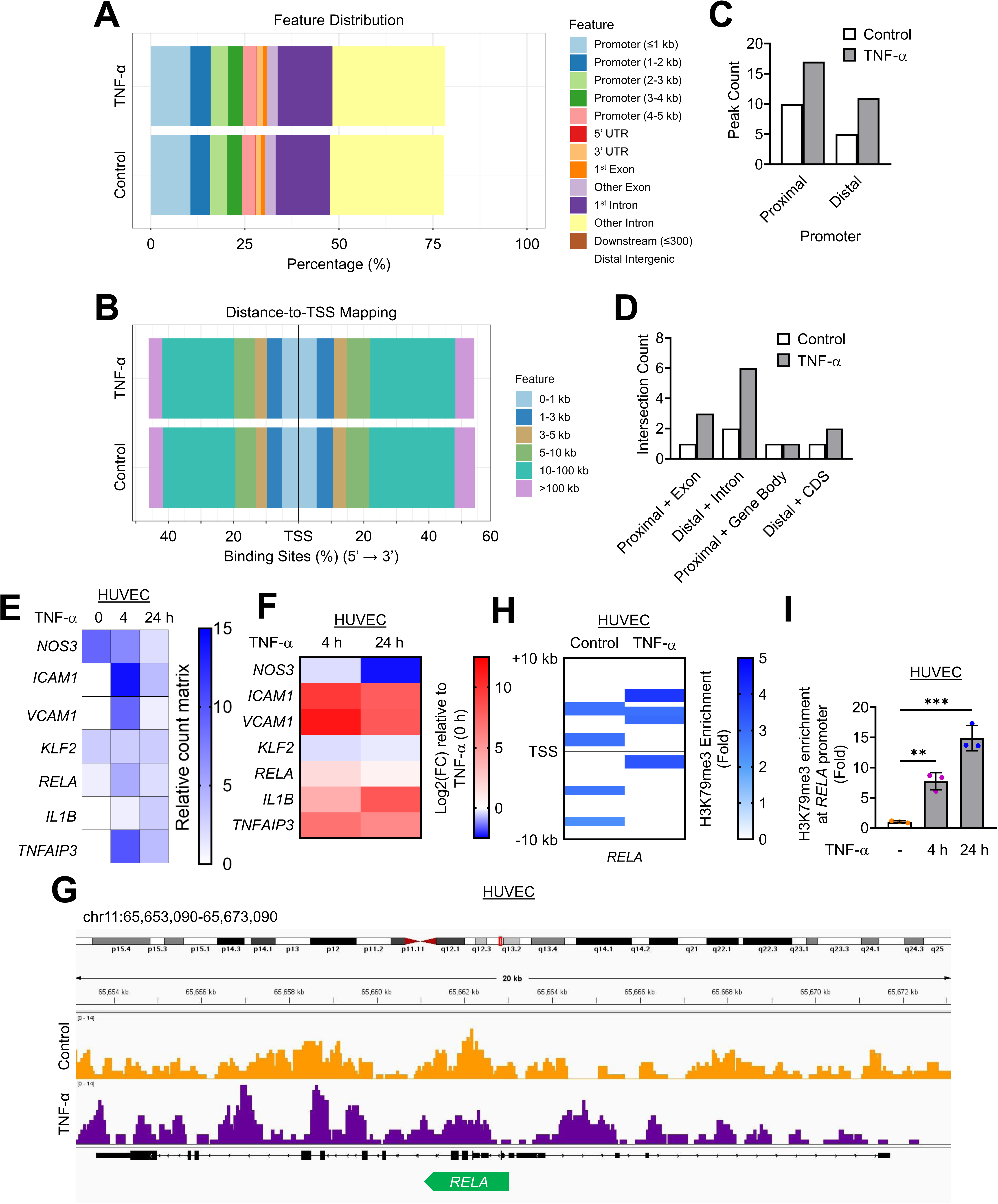
Genome-wide characterization of H3K79me3 distribution and selective enrichment at the *RELA* promoter following TNF-α stimulation in endothelial cells. (A) Genomic feature distribution of H3K79me3 CUT&RUN peaks across promoter-proximal regions, exons, introns, untranslated regions, downstream regions, and distal intergenic loci in control (untreated) and TNF-α–treated (10 ng/ml, 4 h) HUVEC. (B) Distance-to-transcription start site (TSS) analysis showing the positional distribution of H3K79me3 peaks relative to gene promoters (0–1 kb to >100 kb) in untreated and TNF-α–treated (10 ng/ml, 4 h) cells. (C–D) UpSet plot analyses depicting promoter-associated H3K79me3 peaks (C) and their overlap with various genomic features (D) in control and TNF-α treated (10 ng/ml, 4 h) cells. (E-F) Heatmap derived from RNA-seq data (GEO: GSE121958) showing relative count matrix (E) and log₂ fold change (F) in selected endothelial inflammatory genes following TNF-α stimulation (4 and 24 h). (G-H) Integrative Genomics Viewer (IGV) tracks showing normalized H3K79me3 CUT&RUN signal (G) and fold enrichment of annotated H3K79me3 peaks (H) across the *RELA* locus (TSS ± 10 kb) in control and TNF-α–treated (10 ng/ml, 4 h) HUVEC. (I) CUT&RUN-qPCR validation of H3K79me3 enrichment at the *RELA* promoter in HUVEC treated with TNF-α (10 ng/ml) for 0, 4, and 24 h. Data are presented as mean ± SD. *p < 0.05, **p < 0.01, ***p < 0.001 by one-way ANOVA with Tukey’s post hoc test.

To determine whether H3K79me3 directly regulates pro-inflammatory gene expression, we analyzed RNA-seq data from TNF-α-treated HUVECs (0, 4, and 24 h; GSE121958). Multiple inflammatory genes were upregulated *ICAM1*, *VCAM1*, *SELE* (encoding E-Selectin), *RELA* (encoding NF-κB p65), *IL1B* (encoding IL1-β), *IL6*, *CXCL8*, *CXCL10*, *CCL2*, *CXCL1/2*, *PTGS2*, *TNFAIP3*, *STAT1/3*, while *NOS3* (encoding eNOS), *KLF2*, *THBD* were downregulated upon TNF-α stimulation (Figure 3E-F, Supplementary Figure 3D). Integrative analysis of CUT&RUN data revealed selective enrichment of H3K79me3 within ±10 kb of the TSS of *RELA* (encoding NF-κB p65) (Figures 3G-H). These findings were validated by CUT&RUN-qPCR, which showed a time-dependent increase in H3K79me3 enrichment at the *RELA* promoter following 4 h and 24 h of TNF-α treatment, relative to untreated controls (Figure 3I). To further determine whether the observed changes in ICAM1 and VCAM1 expression were mediated by direct promoter regulation via H3K79me3 under TNF-α stimulation, we performed an integrative analysis of CUT&RUN datasets to examine H3K79me3 enrichment within ±10 kb of the TSS of the *ICAM1* and *VCAM1* genes. This analysis did not reveal any significant alterations in H3K79me3 occupancy at these genomic loci following TNF-α treatment (Supplementary Figure 4). Together, these results indicate that TNF-α-induced inflammation reconfigures the H3K79me3 epigenetic landscape, with selective promoter-proximal enrichment at *RELA*, thereby promoting NF-κB p65 transcriptional activation in endothelial cells.

### Both disturbed flow and TNF-α stimulation trigger an NF-κB p65-driven inflammatory phenotype in endothelial cells

Following our observation of increased H3K79me3 enrichment at the *RELA* (NF-κB p65) gene promoter in EC exposed to inflammatory stress, we next aimed to investigate the status of NF-κB p65 in EC subjected to D-Flow and TNF-α stimulation. To confirm the induction of an inflammatory endothelial phenotype, we first assessed the expression of eNOS, ICAM1, and the extent of monocyte adhesion to EC. Our analysis revealed a pronounced decrease in eNOS expression (Supplementary Figure 5A-C) and a significant upregulation of ICAM1 (Supplementary Figure 5D-F) in EC exposed to both D-Flow and TNF-α. Additionally, EC exposed to D-Flow showed increased monocyte adhesion at bifurcation regions (D-Flow zones) of the microchannels compared to S-Flow regions (Supplementary Figure 5G).

Having confirmed the induction of an inflammatory endothelial phenotype, we next evaluated the expression and activation levels of NF-κB p65, its inhibitor IκBα, and the IκBα-inhibiting kinase IKKβ. Our findings indicated elevated NF-κB p65 expression in EC subjected to D-Flow compared to those under S-Flow (Figure 4A-B). Moreover, EC treated with TNF-α displayed increased expression and activation of NF-κB p65 via S536 phosphorylation (Figure 4C-D). Parallel analysis revealed elevated S32 phosphorylation of IκBα, accompanied by a marked downregulation of total IκBα expression, suggesting its dissociation from NF-κB p65 upon activation and subsequent degradation (Figure 4E-F). Finally, we assessed the expression and phosphorylation status of the IκBα inhibitor, IKKβ, in TNF-α-treated EC. This analysis demonstrated increased S176/180 phosphorylation of IKKα/β, while total IKKβ protein levels remained unchanged (Figure 4G-H).

**Figure 4.**
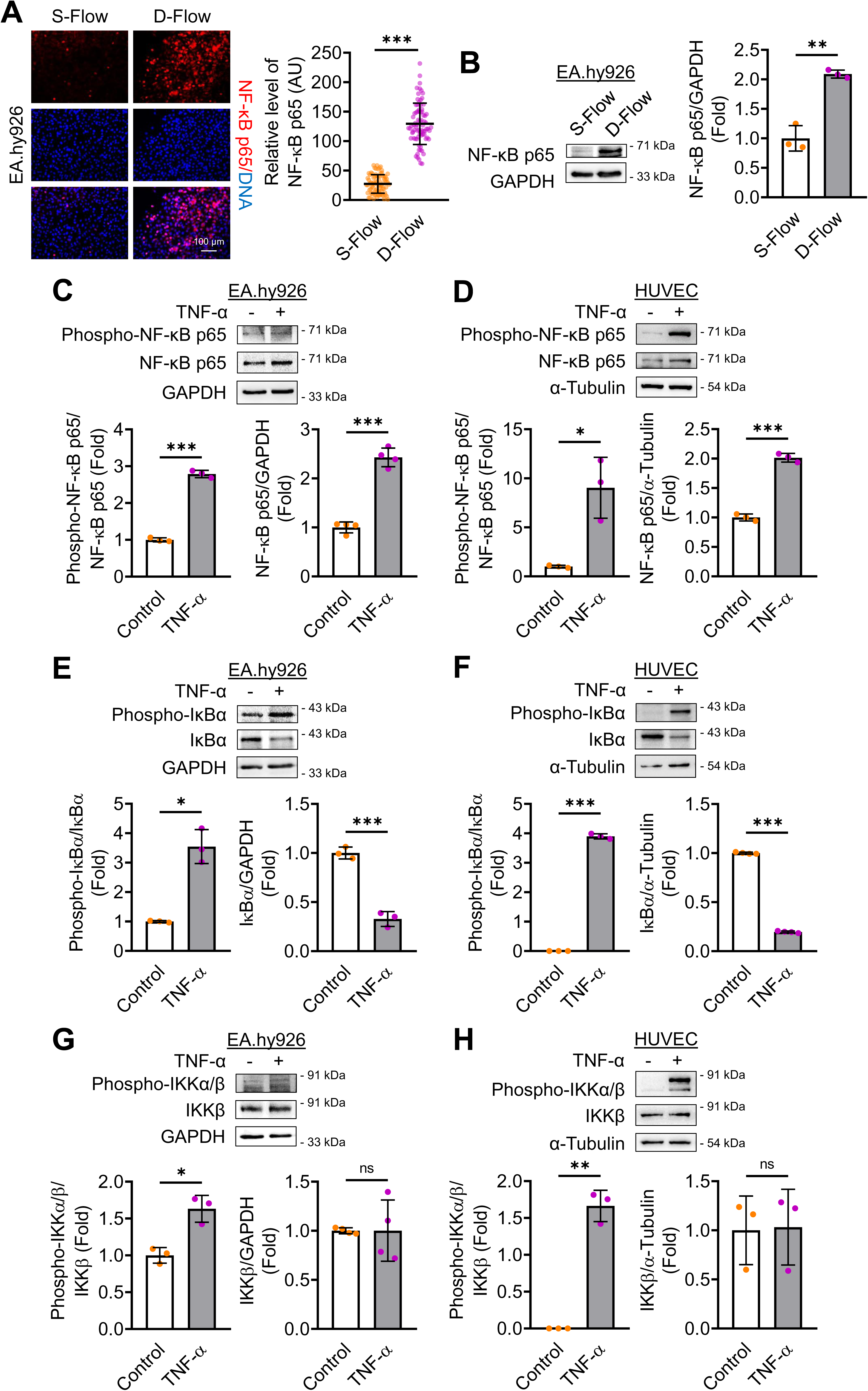
D-Flow and TNF-α stimulation increase NF-κB p65 expression and activation in EC. (A) Immunofluorescence staining of EA.hy926 exposed to D-Flow (4 h; n = 3), showing NF-κB p65 expression (red). Nuclei are stained with DAPI (blue). Fluorescence intensities (AU) in individual EC (dots) from three independent experiments are indicated together with the mean. A total of ≥ 75 EC were analyzed per condition. Magnification: x20; Scale: 100 µm. (B) Immunoblot analysis of NF-κB p65 in EA.hy926 exposed to S- and D-Flow for 4 h (n = 3). (C-D) Immunoblot analysis of Phospho-NF-κB p65 and total NF-κB p65 in EA.hy926 (n = 4; C) and HUVEC (n = 3; D) treated with TNF-α (10 ng/ml, 24 h). (E-F) Immunoblot analysis of Phospho-IκBα and total IκBα in EA.hy926 (n = 4; E) and HUVEC (n = 4; F) following TNF-α stimulation (10 ng/ml, 24 h). (G-H) Immunoblot analysis of Phospho-IKKα/β and total IKKβ in EA.hy926 (n = 4; G) and HUVEC (n = 3; H) treated with TNF-α (10 ng/ml, 24 h). Data are presented as mean ± SD. *p < 0.05, **p < 0.01, ***p < 0.001 by Welch’s t-test.

### Inhibition of DOT1L catalytic activity suppresses NF-κB p65 expression without disrupting other key proteins in the classical NF-κB p65 signaling pathway

Our findings thus far suggest that elevated DOT1L expression enhances H3K79me3 enrichment at the *RELA* (NF-κB p65) gene promoter in EC exposed to both D-Flow and TNF-α. Additionally, we reported increased NF-κB p65 expression and activation in an inflammatory endothelial phenotype. This led us to investigate the role of DOT1L-mediated H3K79me3 deposition at the *RELA* gene promoter in the upregulation and activation of NF-κB p65. To address this, we employed SYC-522, an inhibitor of DOT1L catalytic activity, in EC treated with TNF-α. Our analysis revealed that SYC-522 significantly reduced the level of TNF-α induced H3K79me3 catalysis (Figure 5A, Supplementary Figure 6A). Furthermore, treatment of TNF-α stimulated EC with SYC-522 markedly increased eNOS expression (Figure 5B, Supplementary Figure 6B) while decreasing ICAM1 expression (Figure 5C, Supplementary Figure 6C). Notably, SYC-522 treatment also reduced NF-κB p65 expression and activation via S536 phosphorylation (Figure 5D, Supplementary Figure 6C).

**Figure 5.**
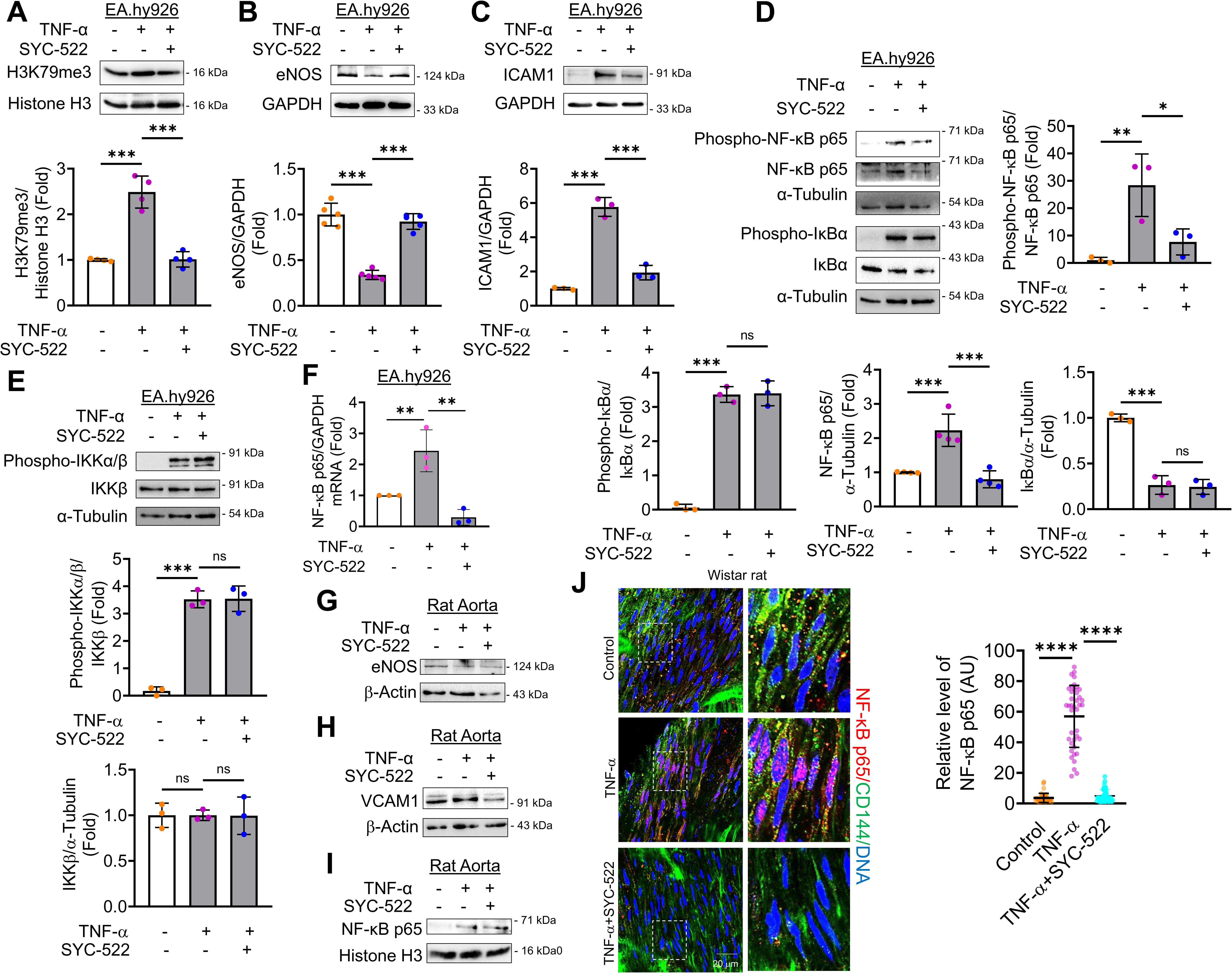
SYC-522 suppresses NF-κB p65 expression and activation independently of its canonical activation in TNF-α treated EC. (A-E) Immunoblot analyses of EA.hy926 treated with TNF-α (10 ng/ml, 24 h) in the presence or absence of the DOT1L inhibitor SYC-522 (5 µM, 24 h), showing the expression of H3K79me3 (n = 4; A), eNOS (n = 5; B), ICAM1 (n = 3; C), Phospho-NF-κB p65 (n = 4), total NF-κB p65 (n = 3), Phospho-IκBα (n = 3), total IκBα (n = 3; D), Phospho-IKKα/β (n = 3), and total IKKβ (n = 3; E). (F) Analyzing the transcript levels of NF-κB p65 in HUVEC EA.hy926 treated with TNF-α (10 ng/ml, 24 h) in the presence or absence of DOT1L siRNA (siDOT1L, 40 nM). (n = 3) (G-I) Immunoblot analyses of rat aortic rings treated with TNF-α (10 ng/ml, 24 h) *ex vivo* in the presence or absence of the DOT1L inhibitor SYC-522 (5 µM, 24 h), showing the expression of eNOS (n = 3; G), VCAM1 (n = 3; H), and total NF-κB p65 (I, n = 3). (J) *En face* preparations were double stained with anti–VE-cadherin (green, EC marker) and an anti-NF-kB p65 antibody (red, C) in TNF-α and/or DOT1L inhibitor (SYC-522) treated aortic tissue. Data are presented as mean ± SD. *p < 0.05, **p < 0.01, ***p < 0.001, ****p < 0.0001 by one-way ANOVA with Tukey’s post hoc test.

To determine whether the reduced levels of phosphorylated NF-κB p65 in SYC-522-treated EC were due to decreased NF-κB p65 expression or disruption of the NF-κB p65 activation signaling cascade, we assessed the expression and activation levels of IκBα. Our results showed no changes in either total or S32 phosphorylated IκBα expression upon SYC-522 treatment in TNF-α-stimulated EC (Figure 5D, Supplementary Figure 6D). Similarly, SYC-522 treatment did not affect the expression or S176/180 phosphorylation of IKKβ or IKKα/β in EC exposed to TNF-α (Figure 5E, Supplementary Figure 6E). We next examined the effect of DOT1L knockdown on TNF-α–induced *NF-κB p65* gene expression in HUVECs. TNF-α stimulation resulted in a significant upregulation of *NF-κB p65* mRNA levels. However, silencing of DOT1L markedly attenuated this induction, leading to a complete reversal of TNF-α–mediated *NF-κB p65* transcript upregulation (Figure 6F). In summary, our findings suggest that SYC-522 reduces H3K79me3 enrichment, thereby suppressing NF-κB p65 expression without affecting the expression or activation of other proteins involved in the canonical NF-κB p65 signaling pathway. Moreover, the inhibition of DOT1L catalytic activity alleviated NF-κB p65-mediated inflammatory responses in TNF-α-treated EC.

**Figure 6.**
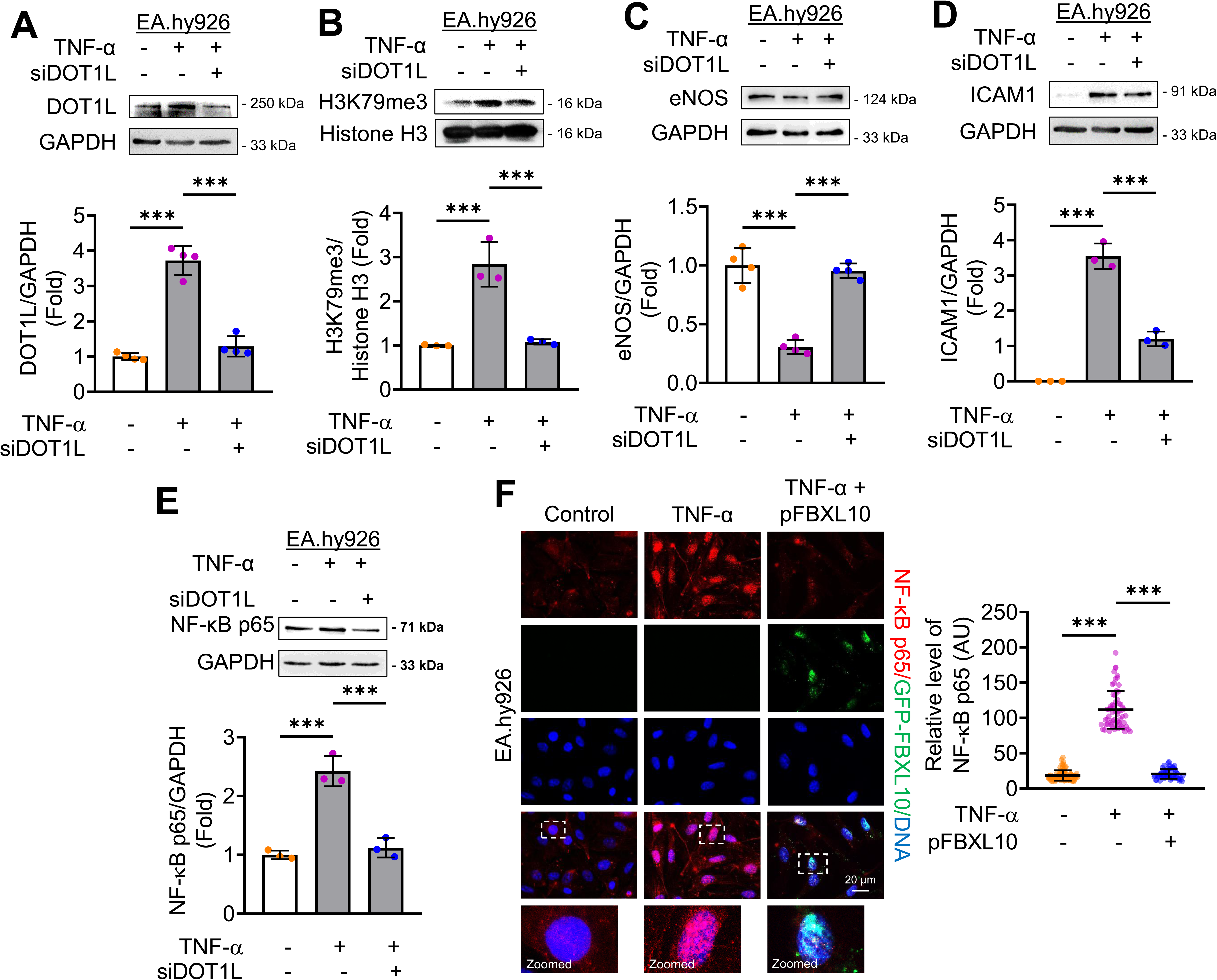
DOT1L silencing and FBXL10 overexpression reduce H3K79me3 enrichment to mitigate NF-κB p65-mediated pro-inflammatory signatures in TNF-α stimulated EC. (A-E) Immunoblot analyses of EA.hy926 treated with TNF-α (10 ng/ml, 24 h) in the presence or absence of DOT1L siRNA (siDOT1L, 40 nM), showing the expression of DOT1L (n = 4; A), H3K79me3 (n = 3; B), eNOS (n = 4; C), ICAM1 (n = 3; D), and NF-κB p65 (n = 3; E). (F) Immunofluorescence staining of EA.hy926 treated with TNF-α (10 ng/ml, 24 h) in the presence or absence of GFP-FBXL10 plasmid (pFBXL10, 250 ng/ml), showing NF-κB p65 expression (red). Nuclei are stained with DAPI (blue). Fluorescence intensities (AU) in individual EC (dots) from three independent experiments are indicated together with the mean. A total of ≥ 55 EC were analyzed per condition. Magnification: x63; Scale: 20 µm. Data are presented as mean ± SD. *p < 0.05, **p < 0.01, ***p < 0.001 by one-way ANOVA with Tukey’s post hoc test.

To further validate our findings in a physiologically relevant setting, we examined the effect of the DOT1L inhibitor SYC-522 in an *ex vivo* model using isolated rat aortic rings exposed to TNF-α. Consistent with our *in vitro* observations, TNF-α treatment resulted in a significant reduction in eNOS protein levels, which was markedly restored upon co-incubation with SYC-522 (Figure 5F). Conversely, TNF-α stimulation significantly increased the expression of the adhesion molecule VCAM1, and this upregulation was attenuated following SYC-522 treatment (Figure 5G). In parallel with the changes observed in VCAM1 expression, TNF-α also elevated total NF-κB p65 protein levels, whereas SYC-522 modestly normalized these alterations (Figure 5G). Furthermore, we also assessed the level of p65 in TNF-α treated rat aortic rings using *en face* staining technique. In parallel with our observation through immunoblot technique, TNF-α also elevated total NF-κB p65 protein levels in endothelial monolayer of the aorta, while SYC-522 normalized the changes in the same (Figure 5J). Collectively, these findings corroborate with our *in vitro* data and support a role for DOT1L in mediating TNF-α induced endothelial inflammatory responses in intact vascular tissue.

### Modulating H3K79me3 enrichment levels restores the NF-κB p65-mediated inflammatory phenotype in endothelial cells

After exploring the potential of pharmacological inhibition of DOT1L in regulating H3K79me3–NF-κB p65-driven endothelial inflammatory signatures, we sought to further investigate the impact of H3K79me3 modulation on the inflammatory endothelial phenotype. To achieve this, we employed a two-pronged approach: siRNA-mediated DOT1L knockdown and FBXL10 overexpression in TNF-α treated EC. FBXL10/KDM2B is a well-characterized demethylase for H3K79me2/3 (23, 24). Our analysis revealed that transient transfection with DOT1L siRNA (siDOT1L) significantly reduced TNF-α-induced DOT1L expression (Figure 6A) and H3K79me3 enrichment (Figure 6B). Moreover, siDOT1L treatment significantly increased eNOS expression (Figure 6C) while reducing ICAM1 (Figure 6D) and NF-κB p65 (Figure 6E) expression levels in TNF-α-treated EC.Additionally, overexpression of the demethylase FBXL10 in TNF-α-treated EC markedly reduced H3K79me3 (Supplementary Figure 7) and nuclear NF-κB p65 expression (Figure 6F).

We then studied the effects of SYC-522 on EC exposed to S-Flow and D-Flow. Our results demonstrated that SYC-522 treatment significantly reduced H3K79me3 enrichment in EC exposed to D-Flow (Figure 7A). Furthermore, SYC-522 reversed the D-Flow-mediated reduction in eNOS expression (Figure 7B) and the increase in NF-κB p65 expression (Figure 7C). Having shown that inhibition of DOT1L-mediated H3K79me3 suppresses D-Flow and TNF-α-induced NF-κB p65 expression in EC, we next assessed the effect of H3K79me3 modulation on NF-κB p65-driven endothelial inflammatory phenotype. To do this, we evaluated the impact of SYC-522 on D-Flow-induced endothelial-monocyte adhesion. Our analysis revealed that SYC-522 treatment significantly reduced the inflammatory characteristics of EC as well as endothelial-monocyte adhesion (Figure 7D). In summary, our findings indicate that D-Flow- or TNF-α-induced DOT1L expression enriches H3K79me3 at the *RELA* (NF-κB p65) gene promoter, thereby regulating NF-κB p65-driven endothelial inflammatory phenotype.

**Figure 7.**
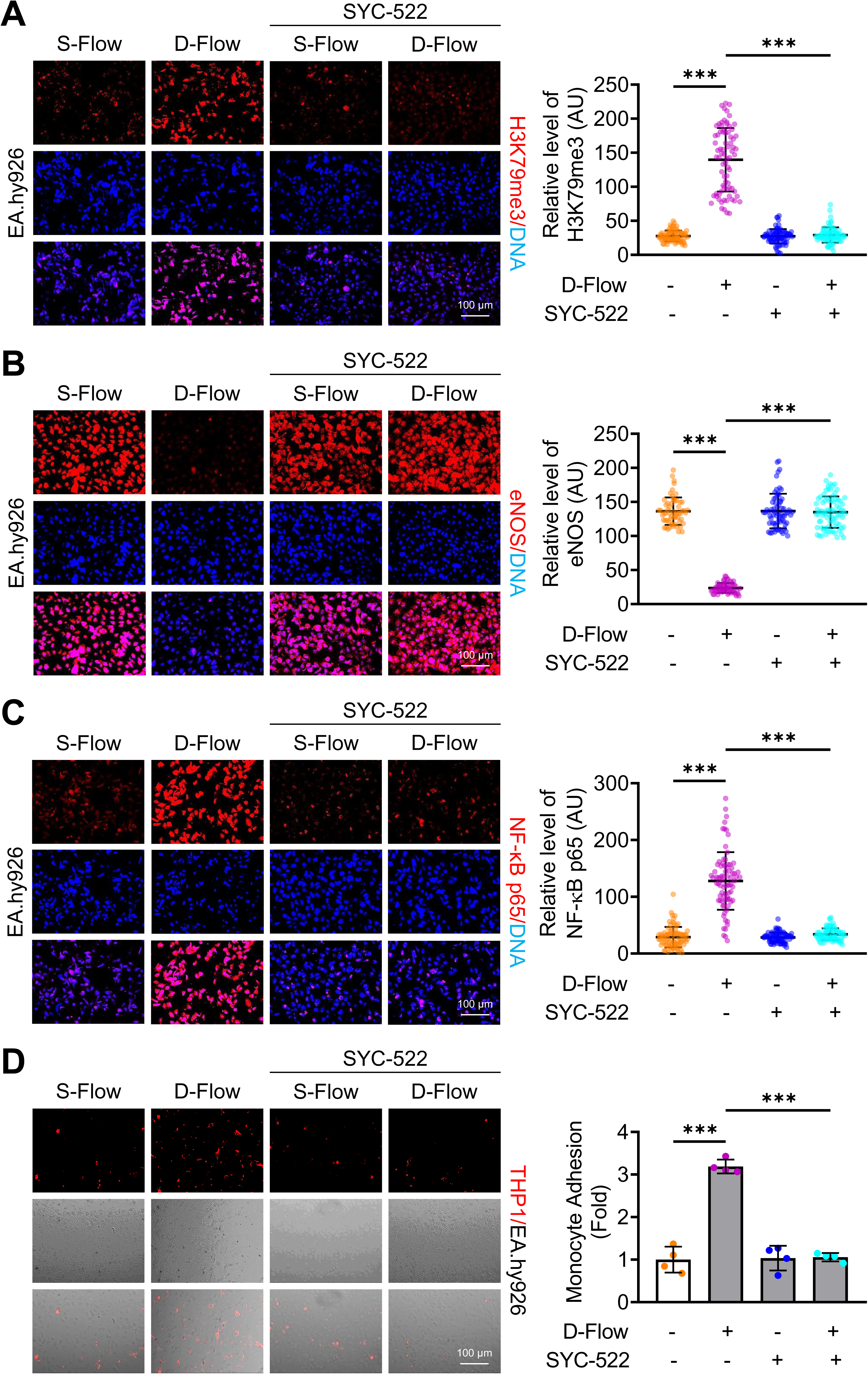
SYC-522 reduces H3K79me3 and attenuates NF-κB p65-driven inflammatory endothelial phenotype under D-Flow conditions. (A-C) Immunofluorescence staining of EA.hy926 exposed to D-Flow (4 h; n = 3), showing the expression of H3K79me3 (A), eNOS (B), and NF-κB p65 (C). Nuclei are stained with DAPI (blue), and target signals corresponding to each respective protein are in red. Fluorescence intensities (AU) in individual EC (dots) from three independent experiments are indicated together with the mean. A total of ≥ 60 EC were analyzed per condition. Magnification: x20; Scale: 100 µm. (D) Monocyte adhesion assay: DiI-labeled THP-1 were incubated (30 min) with flow-exposed EA.hy926 (4 h; n = 4) in the presence or absence of SYC-522 (5 µM). Adherent monocytes (red dots) were counted in regions exposed to S-Flow and D-Flow. Magnification: x10; Scale: 100 µm. Data are presented as mean ± SD. *p < 0.05, **p < 0.01, ***p < 0.001 by one-way ANOVA with Tukey’s post hoc test.

## Discussion

DOT1L-catalyzed methylation of H3K79 is a well-established epigenetic mark associated with transcriptionally active chromatin, playing key roles in development and disease. While inhibition of DOT1L’s catalytic activity is under active investigation, particularly for its anti-cancer potential, its role in pro-atherogenic endothelial inflammation under conditions of disturbed flow remains poorly understood. Here, we show that D-Flow and TNF-α stimulation upregulate DOT1L expression and promote global H3K79me3 enrichment in EC, contributing to a pro-inflammatory phenotype. Specifically, we demonstrated that DOT1L mediates transcriptional priming of the *RELA* (NF-κB p65) gene via H3K79me3 deposition at its promoter, enhancing NF-κB p65 expression and activation. Pharmacological inhibition of DOT1L with SYC-522 reduced both total and phosphorylated NF-κB p65 levels, without altering components of the canonical NF-κB p65 signaling cascade. Moreover, modulating H3K79me3 levels, either by silencing the methyltransferase or by overexpressing the demethylase, reversed the NF-κB p65-driven inflammatory phenotype in EC (Figure 8).

**Figure 8.**
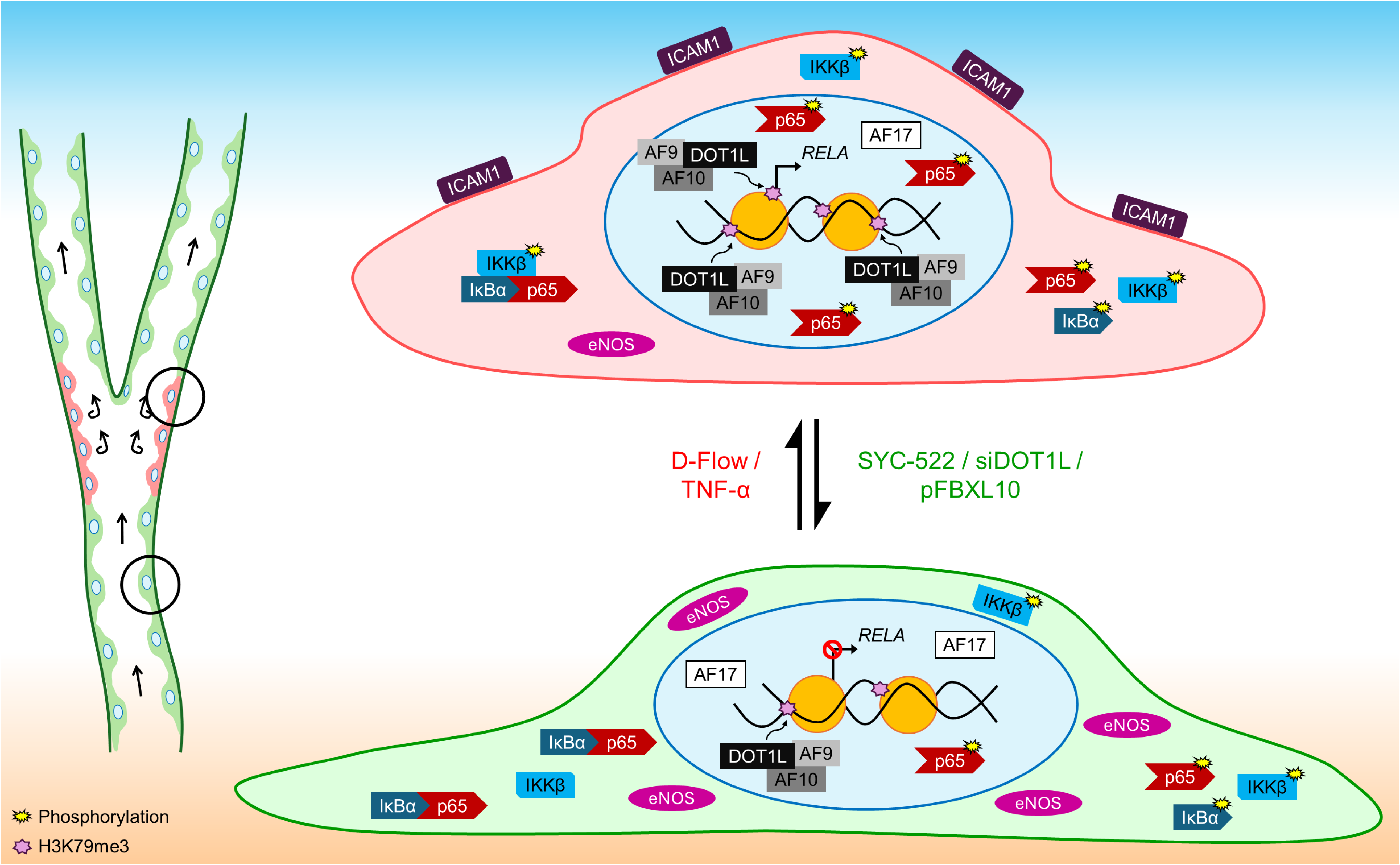
A schematic depicting the critical role of DotCom proteins in endothelial inflammation. D-Flow and TNF-α stimulation enhance DOT1L-catalyzed H3K79me3 enrichment at the *RELA* promoter, thereby promoting a pro-atherogenic inflammatory endothelial phenotype through NF-κB p65. These pro-inflammatory stimuli upregulate the expression of DOT1L and its co-factor/reader AF10, while downregulating AF17, a regulator that facilitates DOT1L nuclear export. This altered DotCom configuration selectively enriches H3K79me3 at the RELA (encoding NF-κB p65) promoter, increasing the pool of NF-κB p65 primed for activation. Although D-Flow and TNF-α activate NF-κB p65 via the canonical pathway, inhibition of DOT1L catalytic activity using SYC-522 suppresses both the expression and activation of NF-κB p65, independent of upstream canonical signaling. Moreover, targeted reduction of H3K79me3–either by DOT1L silencing or FBXL10 overexpression–effectively attenuates NF-κB p65-driven endothelial inflammation.

DOT1L has been a subject of extensive research since its discovery, particularly due to its role in histone methylation and gene regulation. Early studies in *Saccharomyces cerevisiae* identified DOT1L as a histone methyltransferase responsible for H3K79 methylation, demonstrating its role in transcriptional regulation and gene silencing via chromatin modification (25). Subsequent studies uncovered DOT1L’s oncogenic role, demonstrating that it interacts with MLL fusion proteins in leukemic cells, driving aberrant gene expression through dysregulated H3K79 methylation (3). As a result, focus shifted on developing DOT1L inhibitors as potential therapeutic agents. Inhibition of DOT1L was shown to suppress the proliferation of MLL-rearranged leukemia cells (4). These findings spurred the development of DOT1L inhibitors such as EPZ-5676 (Pinometostat), which showed clinical efficacy in MLL-rearranged leukemia patients (26). More recent studies have broadened our understanding of DOT1L’s functions beyond leukemia, revealing its involvement in various developmental processes and diseases. For instance, Jones et al. demonstrated that DOT1L-mediated H3K79 methylation is essential for heterochromatin organization and normal embryonic development. Loss of DOT1L resulted in severe developmental defects, including growth retardation, impaired angiogenesis, and cardiac dilation (5). In addition, studies have linked DOT1L to neural development (6), and neurological and psychotic disorders (8, 27, 28). A study by Cattaneo et al. underscored the critical role of DOT1L-mediated H3K79me2 in cardiomyogenesis, showing that it regulates key transcriptional programs essential for cardiac differentiation (7). Moreover, emerging evidence suggests that DOT1L is also involved in the development and functional maintenance of lymphatic EC (10). The work of Willemsen et al. provided evidence for DOT1L’s role in regulating macrophage lipid biosynthesis, hyperactivation, and the stabilization of atherosclerotic plaques (11), further underscoring the relevance of H3K79 methylation in inflammatory diseases. Another study suggested that DOT1L-driven H3K79me2 regulates the phenotypic transition of VSMC, enhances VSMC-monocyte interactions, and promotes neointimal growth (12), highlighting DOT1L as a promising therapeutic target in vascular pathologies. These studies highlight the critical role of DOT1L-catalyzed H3K79 methylation in VSMC hyperproliferation, neointima formation, and atherosclerotic plaque stabilization. Consequently, we became interested in investigating DOT1L’s role in endothelial dysfunction, a key trigger for the development and progression of atherosclerosis.

We began by assessing DOT1L and H3K79 methylation expression levels *in vivo*, using a D-Flow model by partially ligating the carotid artery, as reported by Nam et al. (15). This revealed increased endothelial expression of DOT1L and H3K79me3, which paralleled neointimal growth and elevated DOT1L expression in VSMC, consistent with previous findings by Farina et al. (12). Next, we validated these *in vivo* observations in an *in vitro* setting. We utilized a 3D-printed model of the human coronary artery with a bifurcation angle of 60°, which effectively replicates *in vivo*-like oscillatory or recirculatory D-Flow patterns (16). In addition, to induce an inflammatory phenotype in EC, we treated them with TNF-α, a well-established method for simulating inflammatory conditions (17). EC exposed to both D-Flow and TNF-α exhibited increased expression of DOT1L and H3K79me3, although we did not observe changes in H3K79me and H3K79me2 levels, contrary to prior reports. DOT1L is part of a protein complex known as DotCom, which includes AF10/MLLT10, AF9/MLLT3, AF17/MLLT6, ENL, Skp1, and TRRAP (29). AF10 enhances DOT1L’s catalytic (18) and recruits DOT1L by recognizing unmodified H3K27 (19). Our findings showed increased AF10 expression in EC exposed to D-Flow and TNF-α, aligning with reports of decreased global H3K27 methylation levels under these conditions (17, 30). AF9 binds acetylated H3K9, recruiting DOT1L to promote H3K79 methylation (20), and we observed elevated AF9 expression in TNF-α-treated EC, consistent with studies linking histone H3 acetylation to endothelial dysfunction (31, 32). Conversely, AF17, which negatively regulates H3K79 methylation by exporting DOT1L from the nucleus (21, 22), was downregulated in EC subjected to D-Flow and TNF-α. Although our findings establish DOT1L activation through upregulation of many DotCom family of genes, the precise molecular mechanisms underlying DOT1L induction and the differential regulation of its associated complex components (AF10, AF9, and AF17) were not comprehensively investigated. Notably, RNA sequencing data suggest that modulation of DotCom components likely occurs at the transcriptional level, a possibility that warrants dedicated future investigation. The primary objective of the present study was to define the downstream functional consequences of DOT1L activation rather than to delineate the regulatory network controlling DotCom gene expression.

Genome-wide profiling revealed that H3K79me3 was enriched at intergenic and intronic regions, suggesting a regulatory role in enhancer activity and alternative splicing. Interestingly, TNF-α stimulation led to increased H3K79me3 enrichment at 5′ and 3′ UTRs, implicating this mark in both transcriptional initiation and post-transcriptional mRNA stability. Promoter-level analysis uncovered a modest yet specific enrichment near the transcription start site (TSS), particularly in the proximal promoter region (<1 kb upstream), indicating selective promoter engagement under inflammatory conditions. Among the inflammation-related genes analyzed, *RELA* emerged as the primary target of H3K79me3 enrichment. CUT&RUN-qPCR validated this selective promoter enrichment, correlating with increased NF-κB p65 transcription following TNF-α exposure. These findings align with and extend emerging evidence that DOT1L plays multifaceted roles in gene regulation beyond its traditional association with transcription elongation. Although historically considered an elongation factor, recent studies have demonstrated that DOT1L is critical for transcription initiation. For instance, loss of DOT1L impairs RNA polymerase II recruitment and pre-initiation complex assembly, while having minimal impact on elongation (33). Moreover, DOT1L’s role is not limited to promoters. H3K79me2/3 marks are also found at active enhancers, contributing to chromatin accessibility and enhancer-promoter communication. These enhancer elements, termed KEE (K79 enhancer elements), are associated with high transcriptional activity and epigenetic plasticity. Loss of DOT1L or its catalytic activity reduces H3K27ac levels and accessibility at KEEs, further implicating DOT1L in enhancer maintenance and epigenetic plasticity (2). Collectively, these data highlight DOT1L as a versatile regulator of gene expression, acting at both promoters and enhancers to coordinate transcriptional responses. In the context of endothelial inflammation, this multifaceted role likely underlies the selective H3K79me3 enrichment at the *RELA* promoter and supports a model in which DOT1L modulates NF-κB p65-dependent transcriptional reprogramming. Notably, the epigenetic landscape surrounding H3K79 methylation is further shaped by opposing chromatin regulators such as FBXL10 (KDM2B), which has been reported to demethylate additional transcription-associated histone marks, including H3K4me3 and H3K36me3. Because these modifications are closely linked to promoter activation and transcriptional elongation, FBXL10 may influence endothelial inflammatory gene expression through coordinated remodeling of multiple chromatin states rather than through H3K79me3 alone (34–36). Although the present study primarily focuses on DOT1L-driven H3K79me3 dynamics, future work will be required to determine whether cross-talk between FBXL10-mediated demethylation and DOT1L-dependent methylation contributes to fine-tuning NF-κB–dependent transcriptional responses during vascular inflammation.

NF-κB p65 is a central regulator of inflammatory responses due to its pivotal role in both the initiation and resolution phases of inflammation (37). It governs a wide spectrum of biological processes, including immune and stress-induced responses, cell proliferation, differentiation, tumorigenesis, apoptosis, and tissue remodelling (38, 39). Aberrant NF-κB p65 activation is linked to various chronic inflammatory diseases, including atherosclerosis (40, 41). Canonical NF-κB p65 signalling is typically activated by pro-inflammatory stimuli such as TNF-α, IL-1β, LPS, and antigens, which engage specific cell surface receptors and initiate downstream signaling cascades that culminate in NF-κB p65 activation. Given our observation of increased DOT1L-catalyzed H3K79me3 enrichment at the *RELA* (NF-κB p65) gene promoter in EC treated with TNF-α, we sought to further elucidate the functional significance of this epigenetic modification. We began by assessing the canonical NF-κB p65 signaling pathway in TNF-α-stimulated EC, observing increased NF-κB p65 expression and activation, marked by S536 phosphorylation. This activation was accompanied by elevated phosphorylation of IKKα/β S176/180. Moreover, increased phosphorylation of IκBα at S32, coupled with decreased total IκBα protein levels, suggested its proteasomal degradation following activation.

Having established enhanced DOT1L-mediated H3K79me3 at the *RELA* promoter, alongside increased NF-κB p65 protein levels and activation, we next examined the impact of inhibiting DOT1L catalytic activity on NF-κB p65 expression and activation. Treatment of TNF-α-stimulated EC with SYC-522 resulted in a significant reduction in H3K79me3, NF-κB p65, and ICAM1 expression while promoting increased eNOS expression. These findings demonstrate that SYC-522 attenuates NF-κB p65-driven inflammatory responses in EC. Our data suggest that the elevated NF-κB p65 protein levels in TNF-α-treated EC are likely driven by increased H3K79me3 enrichment at the *RELA* promoter and that inhibition of this histone modification directly reduces NF-κB p65 expression. Interestingly, much of the research on NF-κB p65 in inflammatory contexts has focused on kinase-mediated activation, with limited attention to its transcriptional regulation (42–45). This prompted us to further investigate the transcriptional control of NF-κB p65. Shu et al. reported increased NF-κB p65 mRNA levels in HUVEC treated with TNF-α and LPS, though their study primarily focused on NF-κB p65’s role in regulating VCAM1 transcription (46). Similarly, Choi et al. demonstrated that nuclear IL-33 transcriptionally regulates NF-κB p65 to induce endothelial activation (47). However, most studies on NF-κB p65-driven endothelial activation have targeted its post-translational activation, rather than its transcript levels (14, 48, 49). For example, Csiszar et al. found that Resveratrol attenuates TNF-α-induced endothelial dysfunction by inhibiting NF-κB p65 activation in coronary artery EC (42), while Pan et al. showed that Resveratrol promotes SIRT1-mediated repression of p38 MAPK and NF-κB p65 activation to protect HUVEC from TNF-α-induced inflammation (50). Additionally, Schubert et al. reported the efficacy of natural antioxidants in inhibiting NF-κB p65 activation in inflammatory endothelial phenotype (51), and Loewe et al. showed that dimethylfumarate effectively mitigates TNF-α-induced endothelial inflammation by selectively blocking the nuclear translocation of activated NF-κB p65 (52).

Upon further investigation, we found that NF-κB p65 plays a key role not only in the initiation of inflammation but also in its resolution and tissue repair. Consistent with this, studies in which NF-κB p65 was inhibited after the initiation phase of inflammation reported prolonged inflammation and delayed tissue repair (53, 54). A CRISPR/Cas9-mediated knockout study by Wang et al. revealed a protective role of NF-κB p65 in maintaining vascular homeostasis (55). Furthermore, Rahman and Fazal highlighted the complexity of NF-κB p65 signalling, suggesting that direct inhibition of IKKα/β or NF-κB p65 may impair both defence and tissue repair mechanisms, thereby underscoring the need for more targeted therapeutic approaches (56). Aligning with this, our data showed that SYC-522-mediated inhibition of DOT1L-catalyzed H3K79me3 significantly reduced both phosphorylated and total NF-κB p65 levels without affecting IκBα and IKKβ activation in TNF-α-treated EC. Notably, SYC-522 reduced NF-κB p65 expression to levels observed in untreated EC, rather than completely abolishing its expression, suggesting a more balanced approach to regulating NF-κB p65-driven endothelial dysfunction.

NF-κB p65 is a well-known transcription factor responsive to hemodynamic forces. In EC, its nuclear translocation and activation are transiently elevated by S-Flow and persistently by D-Flow (57, 58). Once activated, NF-κB p65 drives the transcription of key pro-inflammatory genes, including ICAM1, VCAM1, SELE, and various cytokines, molecules that are critical players in the pathogenesis of atherosclerosis (59). Recent research suggests that transcriptional output of NF-κB p65 is context-dependent, varying with both the nature of the stimulus and the cellular environment (37). For example, the repertoire of NF-κB p65 target genes activated in EC by LPS is distinct from that induced by TNF-α. To determine whether inhibiting DOT1L-catalyzed H3K79me3 could mitigate D-Flow-induced NF-κB p65-driven inflammatory endothelial phenotype, we treated EC exposed to D-Flow with SYC-522. Our results demonstrated decreased H3K79me3 enrichment and NF-κB p65 expression alongside increased eNOS levels. Furthermore, SYC-522 reduced D-Flow-induced endothelial-monocyte interaction and adhesion.

In conclusion, our study reveals that EC exposed to D-Flow and TNF-α exhibit elevated expression of DOT1L, AF10, and AF9, with reduced expression of AF17. This endothelial DotCom configuration promotes H3K79me3 enrichment at the *RELA* (NF-κB p65) gene promoter, leading to increased NF-κB p65 expression. Inhibition of DOT1L-catalyzed H3K79me3 remodelling reverses this effect, reducing NF-κB p65 expression (independent of its activation), rescuing inflammatory signatures, and reducing EC-monocyte interaction/adhesion. Our findings suggest that DOT1L-catalyzed H3K79me3 enrichment primes EC for a pro-atherogenic phenotype under D-Flow and TNF-α exposure, and that epigenetic reprogramming through DOT1L inhibition offers a novel therapeutic strategy to counter NF-κB p65-driven inflammatory responses.

## Experimental procedures

### Animals

Male C57BL/6 mice (aged 6 to 8 weeks) were housed in ventilated cages with *ad libitum* access to standard rodent chow and water. These animals were exposed to a 12 h light and 12 h dark cycle with an ambient temperature of 25°C throughout the study. All the procedures were conducted in accordance with the ARRIVE 2.0 guidelines and were approved by the Institutional Animal Ethics Committee (IAEC) of IIT Kharagpur, India (Protocol number: IE-1/PCS-SMST/2.19).

### Carotid artery ligation

Male C57BL/6 mice (n = 3) were fed a High Fat Rodent Diet (HFD; #D12108C, Research Diets Inc., New Delhi, India) for 10 weeks (6 weeks pre- and 4 weeks post-surgery). Following 6 weeks of HFD feeding, mice were anaesthetized with 1.25% isoflurane (air flow rate: 35 to 50 ml/min) and placed supine on a heating pad. Hair was removed from the region between the mandible and sternum using a depilatory agent, and the area was disinfected with Betadine. A vertical incision (4 – 6 mm) was made in the neck to expose the left carotid artery, identified by blunt dissection adjacent to the trachea. The common carotid was traced caudally to its bifurcation, revealing four branches: the external carotid artery, internal carotid artery, occipital artery, and superior thyroid artery. Partial ligation of the external carotid artery was performed using a 6-0 silk suture (Supplementary Figure 8). The incision was closed with a 4-0 suture and disinfected with Betadine. Postoperatively, a single subcutaneous dose of Meloxicam (2 mg/ml) was administered, and animals were placed in a warm recovery chamber. Four weeks post-surgery, mice were anaesthetized, and blood vessels were perfused and fixed *in situ*. The ligated and contralateral unligated carotid arteries were harvested and cryoprotected in 15 and 30% sucrose solutions. Tissues were embedded in optimal cutting temperature (OCT) compound and stored at -80°C. Transverse sections (7 µm) were obtained using a cryostat (#CM1860, Leica Biosystems, Mumbai, India) and processed for immunostaining.

### *En face* immunohistochemistry

All the experimental procedure involving rodent studies were reviewed and approved by the IAEC of BITS Pilani, Pilani Campus (Protocol Approval No: IAEC/RES/27/12/Rev-1/31/17). We performed this assay as previously described by our group (60). In brief, male Wistar rats aged 12–16 weeks were anesthetized and dissected from the ventral end. PBS was used to perfuse the heart and aorta to eliminate blood cells from the vessels. The primary aortas were then collected, and the fatty tissue layers were removed carefully. For acquiring the aortic explants, the aorta was cut in a size of 8-10 mm cylindrical pieces. Following a PBS wash, the explants were cultured in HiEndoXL™ EC expansion medium and incubated in the complete growth medium for 12 h before initiating the treatment with TNF-α (10 ng/mL) and/or DOT1L inhibitor SYC-522 (5 µM) for 24 h. For analysis of these genes in S-Flow and D-Flow exposed areas of the aorta, specific portion of the aorta from greater curvature experiencing S-Flow and lesser curvature exposed to D-Flow were harvested and processed. Next, aortic rings were cut open longitudinally and permeabilized with PBS containing 0.1% Triton X-100 for 10 min and blocked by TBS containing 10% goat serum and 2.5% Tween 20 for 30 min. Aortas were incubated with 10 µg/ml rabbit anti-DOT1L (#77087, Cell Signaling Technology; rabbit IgG was used as a control) or H3K79me3 (#74073, Cell Signaling Technology; rabbit IgG was used as a control) or NF-κB p65 (#8242, Cell Signaling Technology; rabbit IgG was used as a control) and 12.5 µg/ml mouse anti-VE-cadherin (an EC marker; #sc-9989, Santa Cruz) in the blocking solution overnight. After a PBS rinse, anti–rabbit IgG and anti–mouse IgG (1:1000 dilution; Alexa Fluor 546 and 488, respectively; Invitrogen) were applied for 1 h at room temperature. Nuclei were stained using DAPI (Invitrogen). Images were acquired using a fully Spectral Confocal Laser Scanning Microscope (#LSM 880, Carl Zeiss).

### Cell culture and transient transfection

Human Umbilical Vein Endothelial Cells (HUVEC; #CL002, HiMedia Laboratories, Mumbai, India) were maintained using the Endothelial Cell Expansion Medium (#AL517, HiMedia). EA.hy926 endothelial cells (#CRL-2922, ATCC, Manassas, USA) were grown in DMEM (#AL151A, HiMedia). THP-1 human monocytes (kindly gifted by Prof. Anil Jindal, Department of Pharmacy, BITS Pilani, India) were cultured in RPMI-1640 medium (#AL028A, HiMedia). All cell lines were maintained at 37°C in a humidified incubator with 5% CO_2_.

Human DOT1L siRNA (Sense: 5’-AGAAGCUGUUGAAGGAGAAUU-3’; Antisense: 5′-UUCUCCUUCAACAGCUUCUUU-3′) was procured from GeneCust (Boynes, France) (61). GFP-FBXL10 was a gift from Michele Pagano (Addgene plasmid #126542; http://n2t.net/addgene:126542; RRID: Addgene_126542). For transient transfection, EC were incubated with either DOT1L siRNA or GFP-FBXL10 plasmid using Lipofectamine 2000 Transfection Reagent (Invitrogen, Bengaluru, India) for 4 h. After transfection, cells were cultured in complete medium for a minimum of 36 h prior to treatment with TNF-α (10 ng/ml; 24 h).

### *In vitro* shear/inflammatory stress exposure

PDMS microchannels were fabricated using a 3D-printed human coronary artery mold with a bifurcation angle of 60°. EA.hy926 (6 x 10^5^) were seeded into the channels and incubated for 4 h to allow monolayer formation. DMEM was perfused through the microchannels using the MasterFlex^®^ C/L^®^ Analog Variable-Speed Pump System (Cole-Parmer, Mumbai, India) for 4 h to expose the endothelial monolayer to physiological shear stress. These shear stress-conditioned monolayers were then utilized for immunostaining experiments. For inflammation studies, EA.hy926 or HUVEC were treated with TNF-α (10 ng/ml) for 24 h to induce endothelial activation. In parallel, cells were co-treated with SYC-522 (5 µM) and subsequently processed for immunoblotting.

### RNA Isolation, cDNA Synthesis and qPCR

Reverse transcriptase-quantitative polymerase chain reaction (RT-qPCR) was performed to measure different gene expressions at the transcription level. Upon reaching 80% confluency EA.hy926 cells underwent TNF-α and SYC-522 treatment for 24 h. After that, RNA was isolated from the cells using Trizol Reagent (#15596, TRIzol™ Reagent; Life Technologies, Thermo Fisher Scientific). RNA isolation was succeeded by cDNA preparation from 1 µg of total RNA using iScript™ cDNA Synthesis Kit (#1708891; Bio-Rad Laboratories, Hercules, CA, United States). The quality and quantity of the RNA was measured using a Nano-Drop spectrophotometer (SimpliNano; GE Lifesciences). Before cDNA synthesis, DNA contamination was removed by pre-treating isolated RNAs with the DNase. This was followed by Real-time PCR where iTaq™ Universal SYBR® Green Supermix (#1725124; Bio-Rad Laboratories) was used with a total master mix volume of 10 μl and GAPDH was taken as the housekeeping gene. Data analysis was done by calculating delta-delta Ct. Primer Sequences for *RELA* gene:

Forward: 5’-GCTGCATCCACAGTTTCCAG-3’

Reverse: 5’-TCCCCACGCTGCTCTTCTAT-3’

Primer Sequences for *GAPDH* gene:

Forward Primer: 5’-TCGGAGTCAACGGATTTGGT-3’

Reverse Primer: 5’-TTCCCGTTCTCAGCCTTGAC-3’

### Immunostaining

EC subjected to shear stress or inflammatory stimulation were rinsed with phosphate-buffered saline (PBS) and fixed with 2% paraformaldehyde for 10 min. Cells were permeabilized using 0.1% Triton X-100 and subsequently blocked with 3% bovine serum albumin (BSA) for 1 h at room temperature. For tissue sections, permeabilization was performed using 0.5% Triton X-100, followed by blocking with 10% goat serum supplemented with anti-mouse IgG (1:1000) for 1 h. Cells and tissue sections were incubated overnight at 4°C with the following primary antibodies: DOT1L Rabbit mAb (1:500; #90878), H3K79me3 Rabbit mAb (1:500; #74073), eNOS Rabbit mAb (1:100; #32027), ICAM1 Rabbit pAb (1:500; #4915), NF-κB p65 Rabbit mAb (1:800; #8242, Cell Signaling Technology, Danvers, USA), H3K79me3 Rabbit pAb (1:50; #PA5-96121), AF10 Rabbit pAb (1:500; #BS-3696R), AF17 Rabbit pAb (1:1000; #A302-198A, Invitrogen), and CD144 Mouse mAb (1:25; #SC-9989, Santa Cruz Biotechnology, Dallas, USA). Following three washes with PBS, samples were incubated for 2 h at room temperature with appropriate secondary antibodies: Anti-Rabbit IgG F(ab’)_2_ Fragment Alexa Fluor 555 Conjugate (1:2000; #4413, Cell Signaling Technology), Anti-Rabbit IgG Alexa Fluor 555 Conjugate (1:4000; #A32732,), and/or Anti-Mouse IgG Alexa Fluor 488 Conjugate (1:4000; #A-11001, Invitrogen). Nuclei were counterstained with DAPI for 10 min. Fluorescence imaging was performed using the Zeiss ApoTome 2.0 Microscope (Carl Zeiss), and signal intensities were quantified using ImageJ software.

### Immunoblotting

EA.hy926 and HUVEC were rinsed with PBS and lysed by scraping in RIPA lysis buffer supplemented with 0.1% protease inhibitors. The lysates were sonicated and centrifuged to remove cellular debris. Protein concentrations were determined using the Bradford assay. Equal amounts of total protein were mixed with Laemmli sample buffer and boiled at 100°C for 10 min. Proteins were resolved by SDS-PAGE and transferred onto nitrocellulose membrane. Membranes were blocked with 3% BSA for 1 h at room temperature and incubated overnight at 4°C with the following primary antibodies: DOT1L Rabbit mAb (1:1000; #77087), H3K79me Rabbit mAb (1:1000; #12522), H3K79me2 Rabbit mAb (1:1000; #5427), H3K79me3 Rabbit mAb (1:1000; #74073), eNOS Rabbit mAb (1:1000; #32027), ICAM1 Rabbit pAb (1:1000; #4915), Phospho-NF-κB p65 (Ser536) Rabbit mAb (1:1000; #3033), NF-κB p65 Rabbit mAb (1:1000; #8242), Phospho-IκBα (Ser32) Rabbit mAb (1:1000; #2859), IκBα Mouse mAb (1:1000; #4814), Phospho-IKKα/β (Ser176/180) Rabbit mAb (1:1000; #2697), IKKα Mouse mAb (1:1000; #11930), GAPDH Rabbit mAb (1:2000; #5174), GAPDH Mouse mAb (1:2000; #97166), α-Tubulin Mouse mAb (1:2000; #3873), Histone H3 Rabbit mAb (1:2000; #4499), Histone H3 Mouse mAb (1:2000; #14269, Cell Signaling Technology), AF10 Rabbit pAb (1:500; #BS-3696R), AF9 Rabbit pAb (1:1000; #A300-596A), AF17 Rabbit pAb (1:1000; #A302-198A, Invitrogen). After three washes, membranes were incubated for 1 h at room temperature with secondary antibodies: Anti-Rabbit IgG HRP Conjugate (1:2000; #7074) or Anti-Mouse IgG HRP Conjugate (1:2000; #7076, Cell Signaling Technology). Immunoreactive bands were visualized using Clarity or Clarity Max Western ECL substrates (Bio-Rad, Hercules, USA). Band intensities were quantified by densitometric analysis using ImageJ software.

### Monocyte adhesion assay

DiI stain (1,1’-Dioctadecyl-3,3,3’,3’-Tetramethylindocarbocyanine Perchlorate; #D282, Invitrogen) was generously provided by Prof. Aniruddha Roy (Department of Pharmacy, BITS Pilani, India). THP-1 monocytes (2 x 10^5^ cells per microchannel) were labeled with DiI and incubated with EA.hy926 monolayers previously exposed to *in vitro* fluid shear stress, with or without SYC-522 (5 µM), for 30 min. Non-adherent cells were removed by gentle rinsing with PBS, and the microchannels were fixed with 2% paraformaldehyde. Fluorescence images were acquired using a Zeiss LSM 880 Confocal Microscope. The number of adhered monocytes (visualized as red fluorescent dots) on the endothelial layer was quantified using ImageJ.

### CUT&RUN sequencing, analysis, and qPCR

HUVEC, either untreated or treated with TNF-α (10 ng/ml) for 4 or 24 h, were subjected to Cleavage Under Targets & Release Using Nuclease (CUT&RUN) using the CUT&RUN Assay Kit (#86652, Cell Signaling Technology), following the manufacturer’s instructions. CUT&RUN-seq was performed in two independent experiments (n = 2), wherein each experiment consisted of HUVEC pooled from three biological replicates prior to processing. EC from three independent biological settings were pooled and cross-linked before being bound to Concanavalin A-coated magnetic beads. The cells were then permeabilized with Digitonin and incubated overnight at 4°C with H3K79me3 Rabbit mAb (1:50; #74073, Cell Signaling Technology). Chromatin-bound antibodies were targeted by pAG-MNase, and enzymatic digestion was initiated with calcium chloride. The resulting DNA fragments were incubated at 37°C to release enriched chromatin. Input DNA was prepared in parallel. DNA from both input and CUT&RUN-enriched samples was purified using the DNA Purification Buffers and Spin Columns Kit (#14209, Cell Signaling Technology).

For genome-wide profiling, DNA libraries were constructed from input and H3K79me3-enriched samples using the NEBNext^®^ Ultra™ II DNA Library Preparation Kit (#E7103, New England Biolabs, Gurugram, India) according to the manufacturer’s protocol. High-throughput paired-end sequencing (2 x 150 bp) was performed on an Illumina NovaSeq 6000 platform (v1.5). Raw sequencing data (FASTQ files) underwent quality control, assessing base quality score distribution and average base content per read. Low-quality bases and reads were removed using Trimmomatic v0.39, and high-quality paired-end reads were aligned to the hg38 human reference genome using BWA-MEM v0.7.12. Peak calling was performed with MACS2 v2.1.3 to identify regions of significant H3K79me3 enrichment. Peaks were visualized using the Integrative Genomics Viewer (IGV) and annotated using the ChIPseeker package in R.

To validate genome-wide observations, CUT&RUN-derived DNA from untreated, 4 h, and 24 h TNF-α-treated HUVEC was analyzed by quantitative PCR (qPCR), targeting the *RELA* promoter region. qPCR was performed using the iTaq™ Universal SYBR^®^ Green Supermix (#1725124, Bio-Rad), and fold enrichment was calculated using the ΔΔCt method, with total input DNA serving as the normalization control.

Primer Sequences for *RELA* Promoter:

Forward: 5′-TTCTGTCACCTGAAGTCGGC-3′

Reverse: 5′-GTGGCTGGCCCTGATTAGAA-3′

### RNA sequencing data acquisition, analysis, and visualization

Raw RNA-seq data corresponding to 0, 4, and 24 h TNF-α–treated HUVEC samples were retrieved from the NCBI Gene Expression Omnibus (GEO) under accession number GSE121958. All datasets consisted of paired-end sequencing reads. Quality assessment of the raw reads was performed to ensure data integrity, followed by adapter and low-quality base trimming using Trimmomatic v0.39. The resulting high-quality reads were uploaded to the Galaxy server for downstream processing. Read alignment was done using HISAT2, mapping the sequences to the built-in hg38 human reference genome. Quantification of aligned reads was performed with FeatureCounts, generating read counts mapped to various genomic features including genes, exons, promoters, and genomic bins. These counts were then consolidated into a unified count matrix and downloaded for local analysis. To enable gene-level interpretation, annotation of the gene IDs in the matrix was performed using the AnnotateMyIDs tool. This yielded a tabular file containing Entrez IDs, Ensembl IDs, gene names, and gene functions. Based on this annotation, genes of interest were filtered, and manual calculations were performed to compute fold change and log2 fold change (Log2FC) values. The processed dataset containing Log2FC values was imported into Prism, where heatmaps were generated to visualize gene expression dynamics across the different TNF-α treatment conditions.

### Statistics

All data are presented as mean ± SD. Statistical differences between two groups were assessed using Welch’s *t*-test, while comparisons among three or more groups were evaluated using one-way ANOVA followed by Tukey’s post-hoc test. Statistical analyses were performed using GraphPad Prism versions 9.5.1 and 10. A *p*-value < 0.05 was considered statistically significant. Unless otherwise mentioned, all experiments were conducted with a minimum of three independent biological experimental replicates (n = 3). For immunostaining analyses, quantification was based on multiple cells imaged from distinct fields of view across at least three independent biological replicates.

## Data availability

The raw sequencing files in fastq format and the summarized genome browser tracks in bed format are available in the GEO database. Raw RNA-seq data corresponding to 0, 4, and 24 h TNF-α–treated samples were obtained from the NCBI Gene Expression Omnibus (GEO) under accession number GSE121958 (https://www.ncbi.nlm.nih.gov/geo/query/acc.cgi?acc=GSM3450494). All other raw data can be obtained from the author upon a reasonable request.

## Acknowledgments

The authors gratefully acknowledge the technical assistance of Mr. Suman Kumar in operating the Confocal microscope at the Sophisticated Instrumentation Facility (SIF), BITS Pilani, Pilani Campus, India. We also extend our sincere thanks to Dr. Lakshmikirupa Sundaresan (The Hospital for Sick Children [SickKids], Toronto, Canada) and Dr. Soumitra Ghosh (Department of Biological Sciences, BITS Pilani, Pilani Campus, India) for their valuable support with the downstream analysis of the CUT&RUN sequencing data.

## Author contributions

YTK designed and performed the experiments, analyzed the data, and wrote the initial manuscript draft. RB, SD, SCB, NS, and NG contributed to the experimental work and data analysis. NS and SC conducted the bioinformatic analysis of the CUT&RUN sequencing data. PCS contributed to the experimental design and revised the final manuscript. SM secured the funding, conceived and supervised the study, and wrote and edited the final manuscript.

