## Supplemental Figures for "DOT1L-AF10–mediated H3K79me3 promotes NF-κB p65–dependent inflammatory activation in endothelial cells"

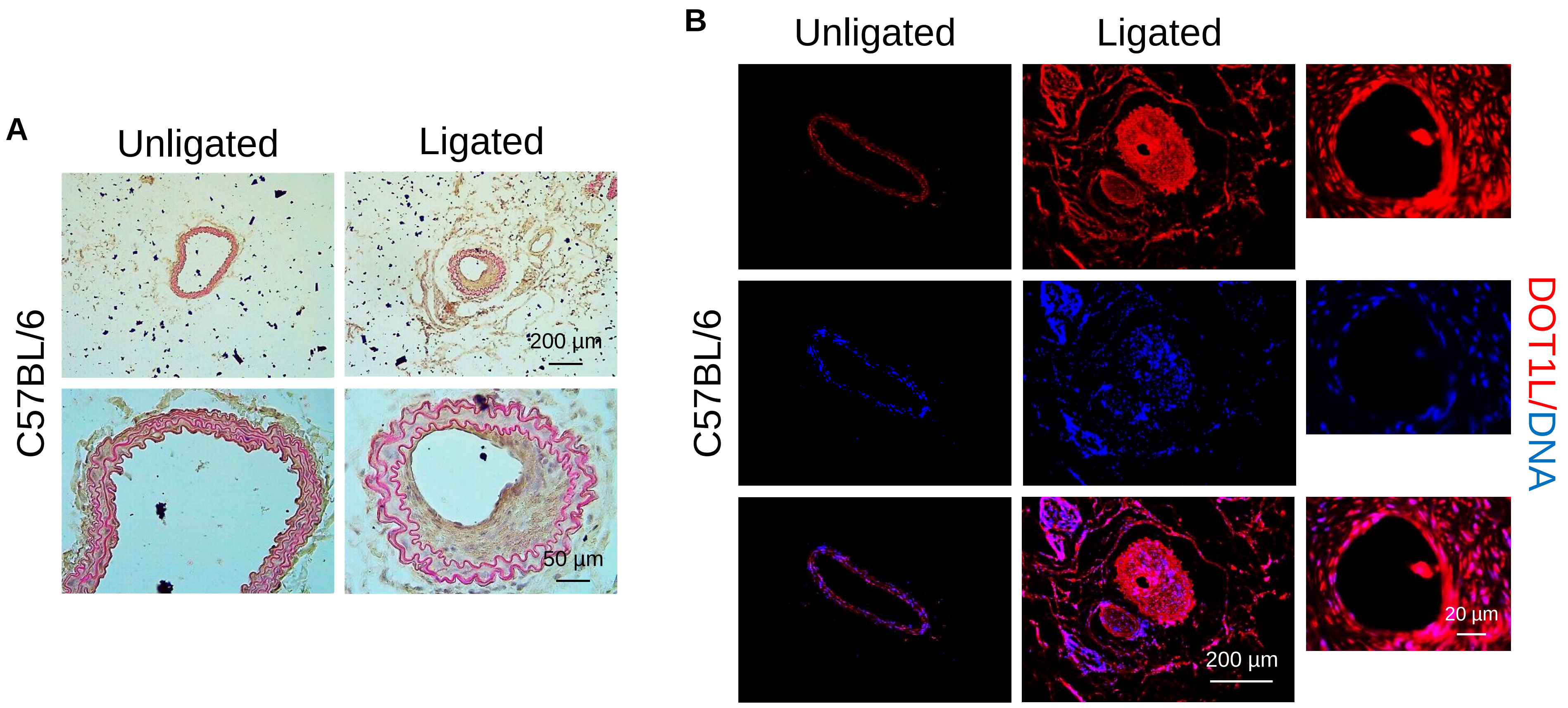


**Supplementary Figure 1. Partial carotid artery ligation induces neointimal hyperplasia and upregulates DOT1L expression *in vivo*.** (A) Hematoxylin and eosin (H&E) staining of carotid artery cross-sections from HFD-fed C57BL/6 mice (n = 3), collected four weeks post-surgery, demonstrating neointimal thickening in the ligated artery compared with the unligated control. Magnification: x10, x40; Scale: 200 µm, 50 µm. (B) Immunofluorescence staining for DOT1L (red) in histological sections of partially ligated and unligated carotid arteries from HFD-fed C57BL/6 mice (n = 3), showing increased DOT1L signal with the neointimal region following ligation. Nuclei are stained with DAPI (blue). Magnification: x10, x63; Scale: 200 µm, 20 µm.

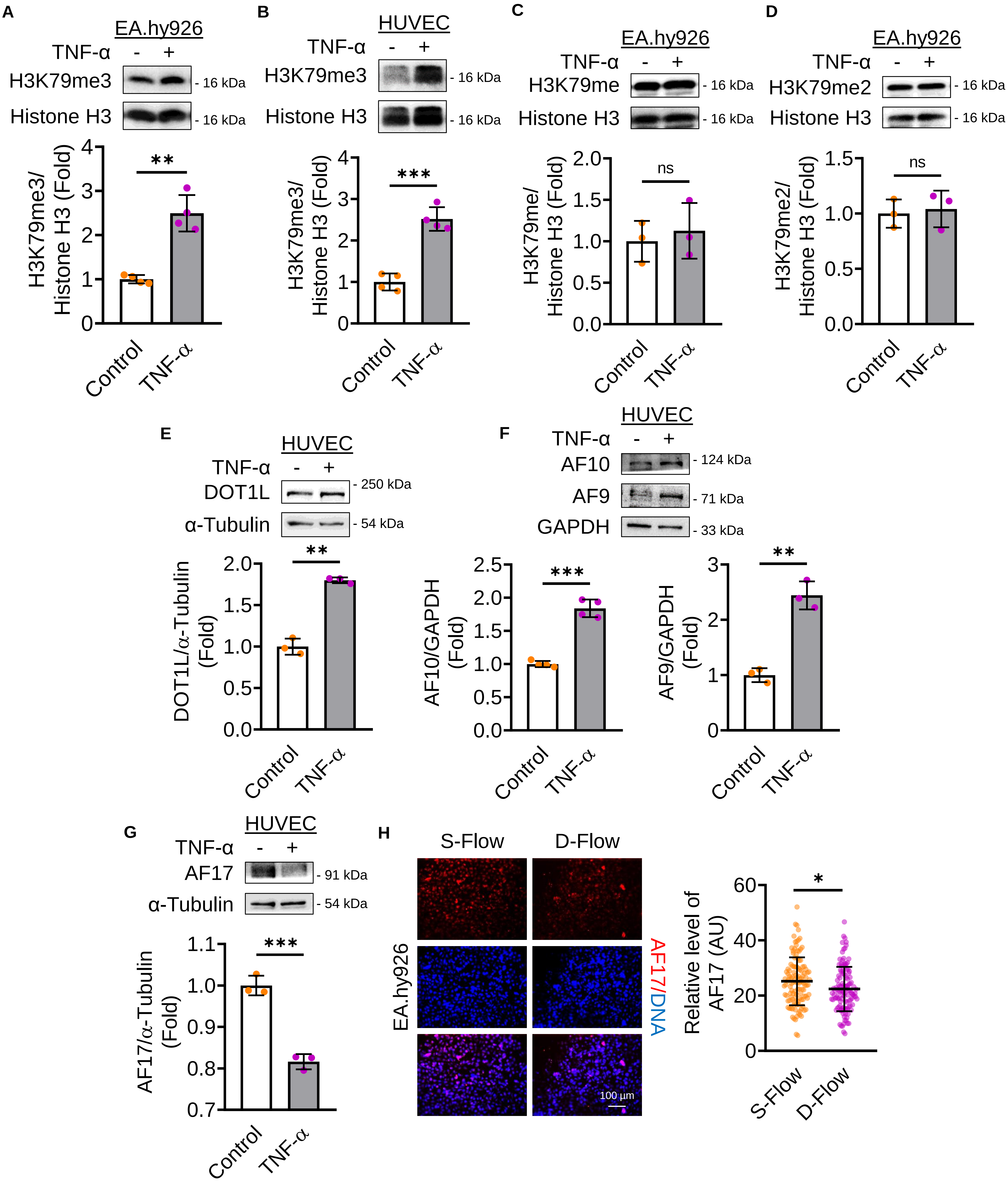
A
EA.hy926
TNF-α - +
H3K79me3
- 16 kDa
Histone H3
- 16 kDa
H3K79me3/
Histone H3 (Fold)
**
Control
TNF-α
B
HUVEC
TNF-α - +
H3K79me3
- 16 kDa
Histone H3
- 16 kDa
H3K79me3/
Histone H3 (Fold)
***
Control
TNF-α
C
EA.hy926
TNF-α - +
H3K79me
- 16 kDa
Histone H3
- 16 kDa
H3K79me/
Histone H3 (Fold)
ns
Control
TNF-α
D
EA.hy926
TNF-α - +
H3K79me2
- 16 kDa
Histone H3
- 16 kDa
H3K79me2/
Histone H3 (Fold)
ns
Control
TNF-α
E
HUVEC
TNF-α - +
DOT1L
- 250 kDa
α-Tubulin
- 54 kDa
DOT1L/α-Tubulin
(Fold)
**
Control
TNF-α
F
HUVEC
TNF-α - +
AF10
- 124 kDa
AF9
- 71 kDa
GAPDH
- 33 kDa
AF10/GAPDH
(Fold)
***
AF9/GAPDH
(Fold)
**
Control
TNF-α
G
HUVEC
TNF-α - +
AF17
- 91 kDa
α-Tubulin
- 54 kDa
AF17/α-Tubulin
(Fold)
***
Control
TNF-α
H
S-Flow
D-Flow
EA.hy926
AF17/DNA
100 µm
Relative level of
AF17 (AU)
*
S-Flow
D-Flow

**Supplementary Figure 2. Altered DotCom protein expression selectively enhances H3K79me3 in pro-inflammatory EC.** (A-B) Immunoblot analysis of H3K79me3 levels in EA.hy926 (n = 4; A) and HUVEC (n = 4; B) treated with TNF-α (10 ng/ml, 24 h). (C-D) Immunoblot analysis of EA.hy926 treated with TNF-α (10 ng/ml, 24 h; n = 3), showing H3K79me (C) and H3K79me2 (D) levels. (E-G) Immunoblot analyses of HUVEC treated with TNF-α (10 ng/ml, 24 h), showing the expression of DOT1L (n = 3; E), AF10 (n = 4), AF9 (n = 3; F), and AF17 (n = 3; G). (H) Immunofluorescence staining of EA.hy926 exposed to D-Flow (4 h; n = 3), showing AF17 expression (red). Nuclei are stained with DAPI (blue). Fluorescence intensities (AU) in individual EC (dots) from three independent experiments are indicated together with the mean. A total of ≥ 120 EC were analyzed per condition. Magnification: x20; Scale: 100 µm. Data are presented as mean ± SD. $^{*}p < 0.05$, $^{**}p < 0.01$, $^{***}p < 0.001$ by Welch’s t-test.

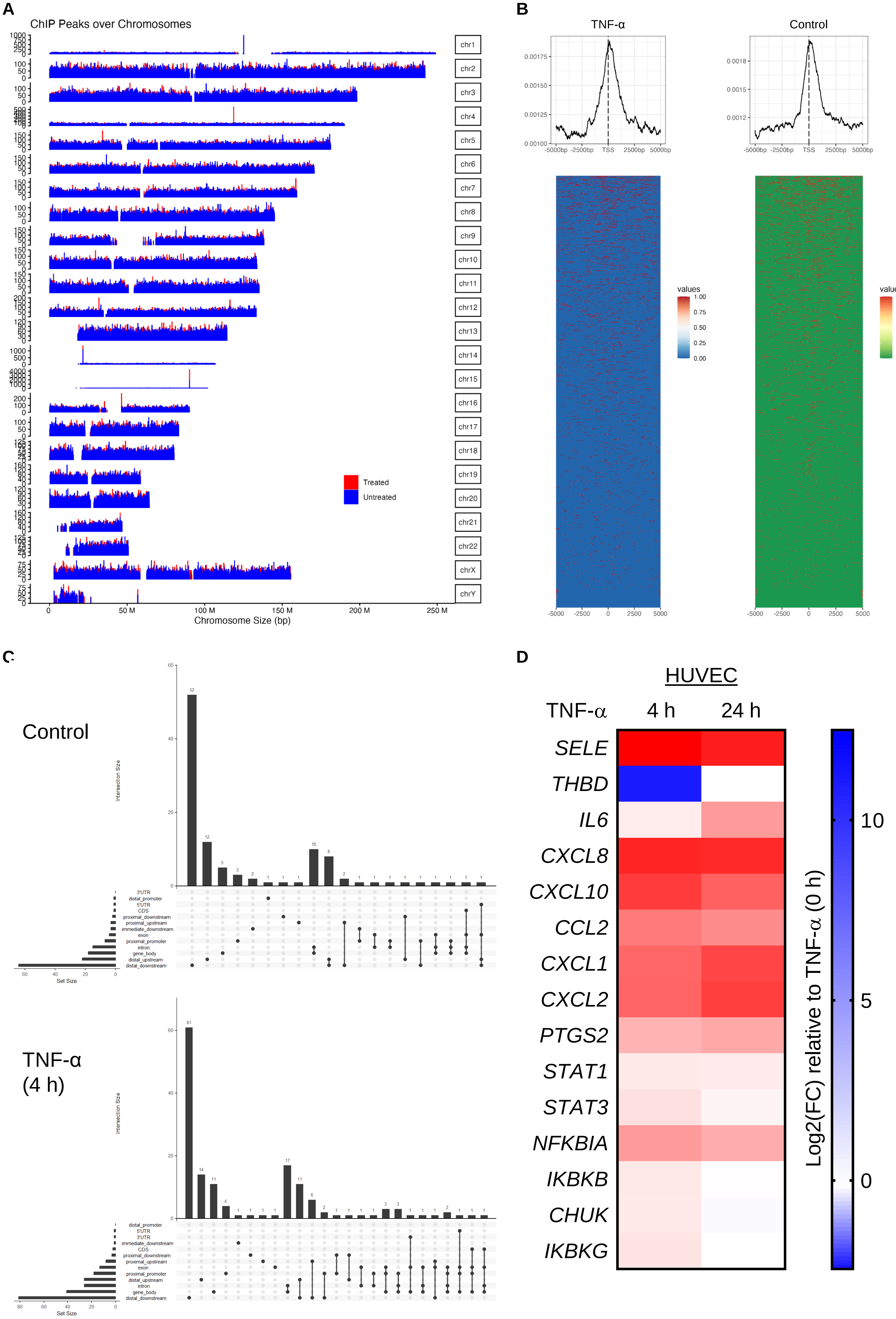
A
ChIP Peaks over Chromosomes
chr1
chr2
chr3
chr4
chr5
chr6
chr7
chr8
chr9
chr10
chr11
chr12
chr13
chr14
chr15
chr16
chr17
chr18
chr19
chr20
chr21
chr22
chrX
chrY
Treated
Untreated
0
50 M
100 M
150 M
200 M
250 M
Chromosome Size (bp)
B
TNF-α
Control
-5000bp -2500bp TSS 2500bp 5000bp
values
1.00
0.75
0.50
0.25
0.00
-5000 -2500 0 2500 5000
C
Control
TNF-α
(4 h)
Intersection Size
Set Size
3'UTR
distal_promoter
5'UTR
CDS
proximal_downstream
proximal_upstream
immediate_downstream
exon
proximal_promoter
intron
gene_body
distal_upstream
distal_downstream
D
HUVEC
TNF-α
4 h
24 h
SELE
THBD
IL6
CXCL8
CXCL10
CCL2
CXCL1
CXCL2
PTGS2
STAT1
STAT3
NFKBIA
IKBKB
CHUK
IKBKG
10
5
0
Log2(FC) relative to TNF-α (0 h)

**Supplementary Figure 3. Genome-wide distribution and functional association of H3K79me3 in TNF-α–stimulated endothelial cells.** (A) Genome-wide tracks showing normalized H3K79me3 CUT&RUN signal across human chromosomes in control (untreated; blue) and TNF-α–treated (10 ng/ml, 4 h; red) HUVEC. (B) Metagene profiles and heatmaps of normalized H3K79me3 enrichment centered on transcription start sites (TSS; −5 kb to +5 kb) in control and TNF-α–treated (10 ng/ml, 4 h) cells. (C) UpSet plot analysis showing genomic feature annotation and combinatorial peak intersections in control and TNF-α–treated (10 ng/ml, 4 h) HUVEC. (D) Heatmap of RNA-seq–derived expression changes for selected inflammatory and endothelial genes.

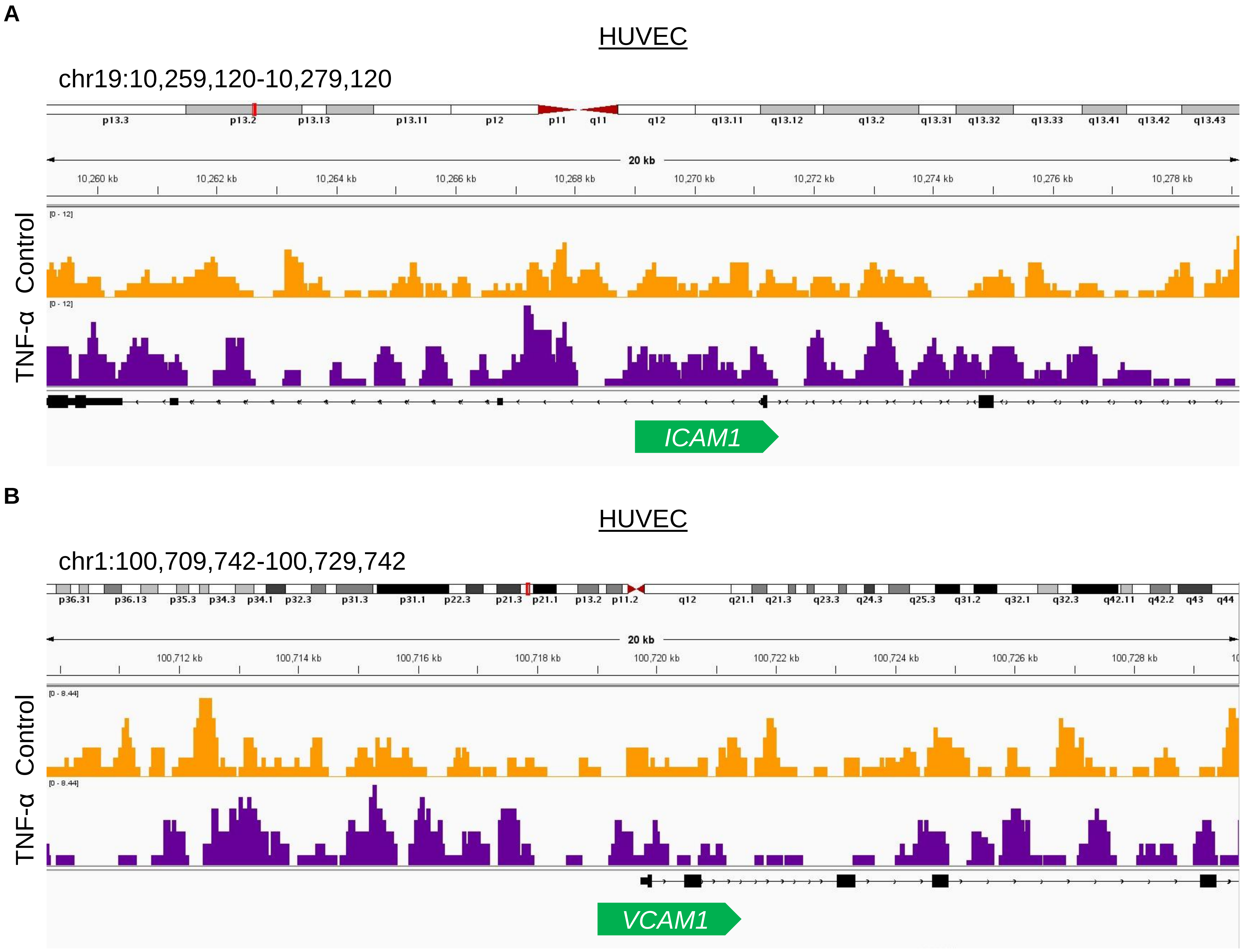


**Supplementary Figure 4. Genome browser visualization of H3K79me3 enrichment at inflammatory endothelial gene loci.** (A-B) Integrative Genomics Viewer (IGV) tracks showing normalized H3K79me3 CUT&RUN signal across the *ICAM1* (A) and *VCAM1* (B) loci (TSS ± 10 kb) in control and TNF-α–treated (10 ng/ml, 4 h) HUVEC.

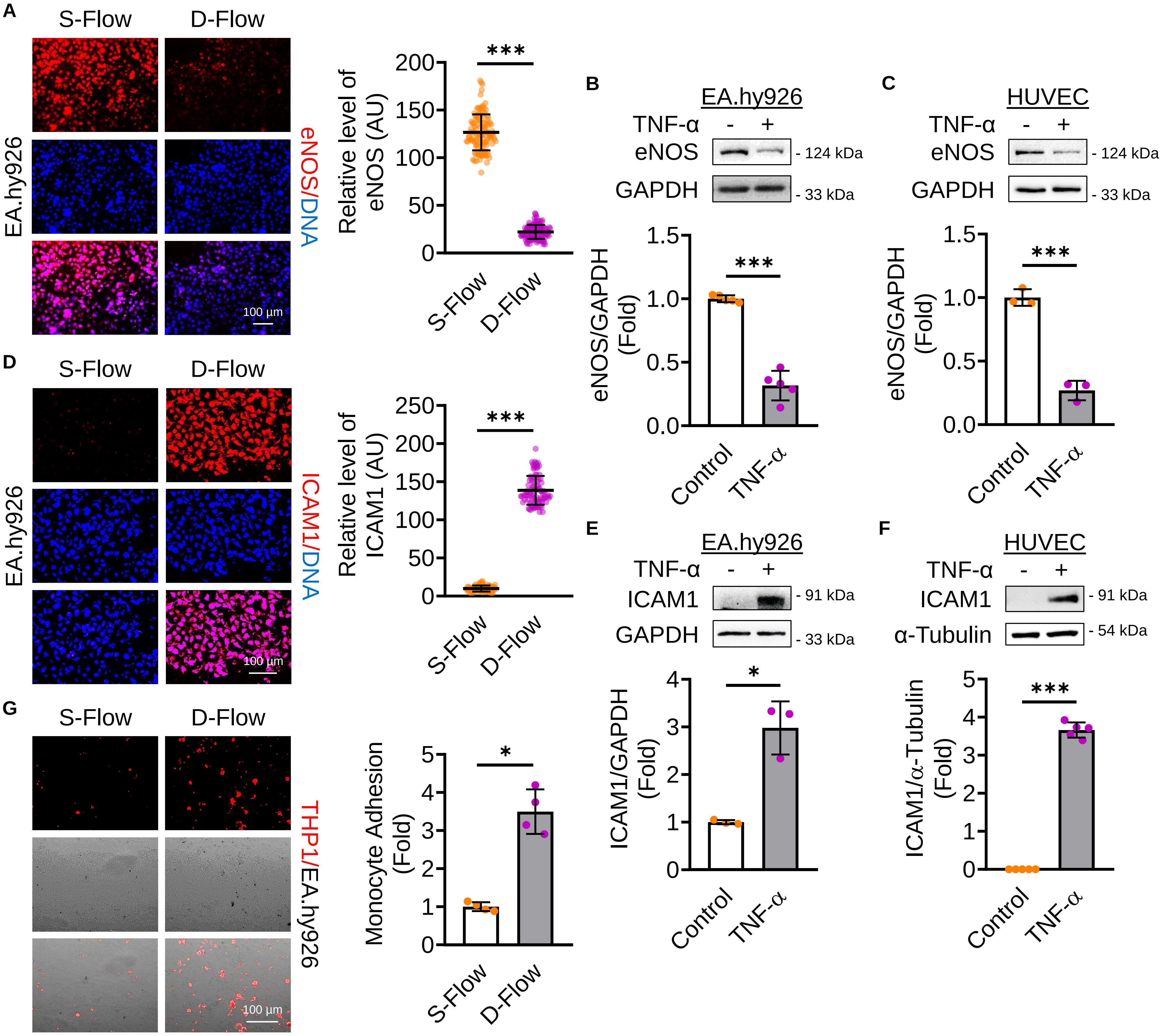
A
S-Flow
D-Flow
EA.hy926
eNOS/DNA
100 µm
Relative level of eNOS (AU)
***
B
EA.hy926
TNF-α - +
eNOS - 124 kDa
GAPDH - 33 kDa
eNOS/GAPDH (Fold)
***
Control
TNF-α
C
HUVEC
TNF-α - +
eNOS - 124 kDa
GAPDH - 33 kDa
eNOS/GAPDH (Fold)
***
D
ICAM1/DNA
Relative level of ICAM1 (AU)
E
ICAM1 - 91 kDa
ICAM1/GAPDH (Fold)
*
F
ICAM1 - 91 kDa
α-Tubulin - 54 kDa
ICAM1/α-Tubulin (Fold)
G
THP1/EA.hy926
Monocyte Adhesion (Fold)

**Supplementary Figure 5. D-Flow and TNF-α exposures induce a pro-inflammatory endothelial phenotype.** (A) Immunofluorescence staining of EA.hy926 exposed to D-Flow (4 h; n = 3), showing eNOS expression (red). Nuclei are stained with DAPI (blue). Fluorescence intensities (AU) in individual EC (dots) from three independent experiments are indicated together with the mean. A total of ≥ 120 EC were analyzed per condition. Magnification: x20; Scale: 100 µm. (B-C) Immunoblot analysis of eNOS expression in EA.hy926 (n = 5; B) and HUVEC (n = 3; C) treated with TNF-α (10 ng/ml, 24 h). (D) Immunofluorescence staining of EA.hy926 exposed to D-Flow (4 h; n = 3), showing ICAM1 expression (red). Nuclei are stained with DAPI (blue). Fluorescence intensities (AU) in individual EC (dots) from three independent experiments are indicated together with the mean. A total of ≥ 75 EC were analyzed per condition. Magnification: x20; Scale: 100 µm. (E-F) Immunoblot analysis of ICAM1 expression in EA.hy926 (n = 3; E) and HUVEC (n = 5; F) treated with TNF-α (10 ng/ml, 24 h). (G) Monocyte adhesion assay: DiI-labelled THP-1 cells were incubated (30 min) with flow-exposed EA.hy926 (4 h; n = 4), and adherent monocytes (red dots) were counted in regions exposed to S-Flow and D-Flow. Magnification: x10; Scale: 100 µm. Data are presented as mean ± SD. *$p < 0.05$, **$p < 0.01$, ***$p < 0.001$ by Welch's t-test.

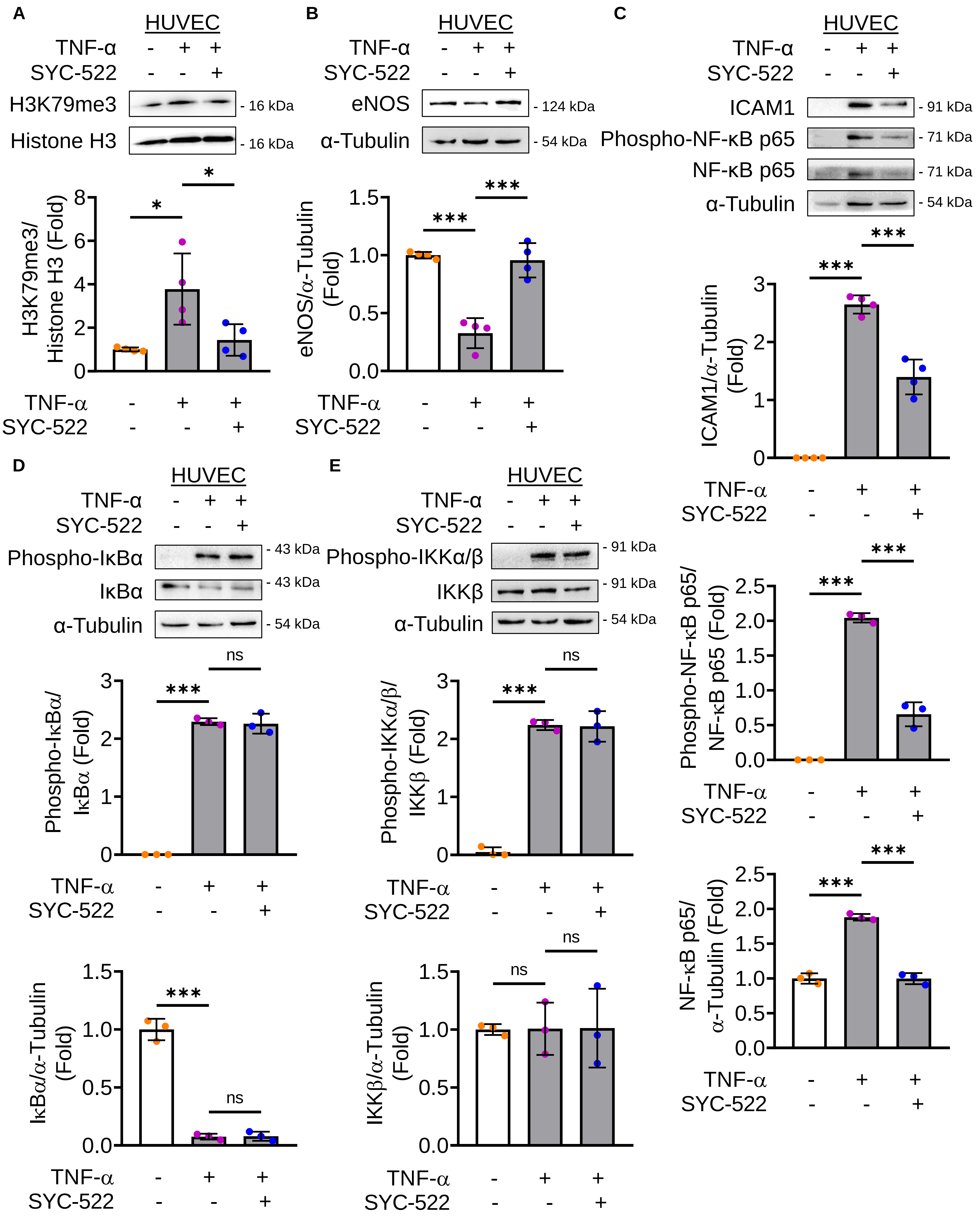

A
HUVEC
TNF-α - + +
SYC-522 - - +
H3K79me3 - 16 kDa
Histone H3 - 16 kDa
H3K79me3/ Histone H3 (Fold)
B
HUVEC
TNF-α - + +
SYC-522 - - +
eNOS - 124 kDa
α-Tubulin - 54 kDa
eNOS/α-Tubulin (Fold)
C
HUVEC
TNF-α - + +
SYC-522 - - +
ICAM1 - 91 kDa
Phospho-NF-κB p65 - 71 kDa
NF-κB p65 - 71 kDa
α-Tubulin - 54 kDa
ICAM1/α-Tubulin (Fold)
Phospho-NF-κB p65/ NF-κB p65 (Fold)
NF-κB p65/ α-Tubulin (Fold)
D
HUVEC
TNF-α - + +
SYC-522 - - +
Phospho-IκBα - 43 kDa
IκBα - 43 kDa
α-Tubulin - 54 kDa
Phospho-IκBα/ IκBα (Fold)
IκBα/α-Tubulin (Fold)
E
HUVEC
TNF-α - + +
SYC-522 - - +
Phospho-IKKα/β - 91 kDa
IKKβ - 91 kDa
α-Tubulin - 54 kDa
Phospho-IKKα/β/ IKKβ (Fold)
IKKβ/α-Tubulin (Fold)
ns
*
***

**Supplementary Figure 6. SYC-522 attenuates TNF-α-induced NF-κB p65 expression and activation in HUVEC independently of the canonical activation pathway.** (A-E) Immunoblot analyses of EA.hy926 treated with TNF-α (10 ng/ml, 24 h) in the presence or absence of the DOT1L inhibitor SYC-522 (5 µM, 24 h), showing the expression of H3K79me3 ($n = 4$; A), eNOS ($n = 4$; B), ICAM1 ($n = 4$), Phospho-NF-κB p65 ($n = 3$), total NF-κB p65 ($n = 3$; C), Phospho-IκBα ($n = 3$), total IκBα ($n = 3$; D), Phospho-IKKα/β ($n = 3$), and total IKKβ ($n = 3$; E). Data are presented as mean ± SD. *$p < 0.05$, **$p < 0.01$, ***$p < 0.001$ by one-way ANOVA with Tukey's post hoc test.

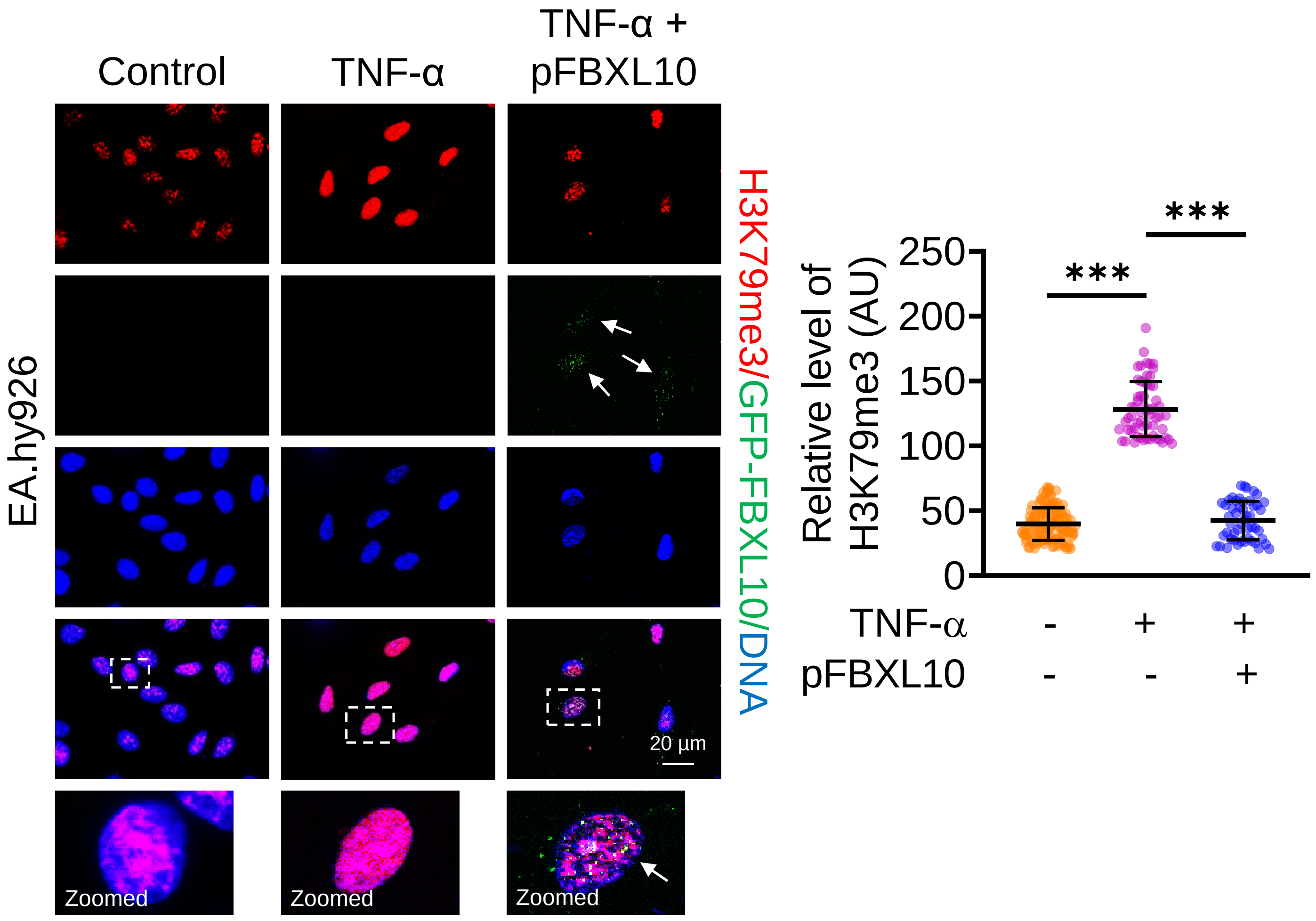


**Supplementary Figure 7. FBXL10 overexpression reduces H3K79me3 enrichment in TNF-α stimulated EC.** Immunofluorescence staining of EA.hy926 treated with TNF-α (10 ng/ml, 24 h) in the presence or absence of GFP-FBXL10 plasmid (pFBXL10, 250 ng/ml), showing H3K79me3 levels (red). Nuclei are stained with DAPI (blue). Fluorescence intensities (AU) in individual EC (dots) from three independent experiments are indicated together with the mean. A total of ≥ 50 EC were analyzed per condition. Magnification: x63; Scale: 20 µm. Data are presented as mean ± SD. *p < 0.05, **p < 0.01, ***p < 0.001 by one-way ANOVA with Tukey's post hoc test.

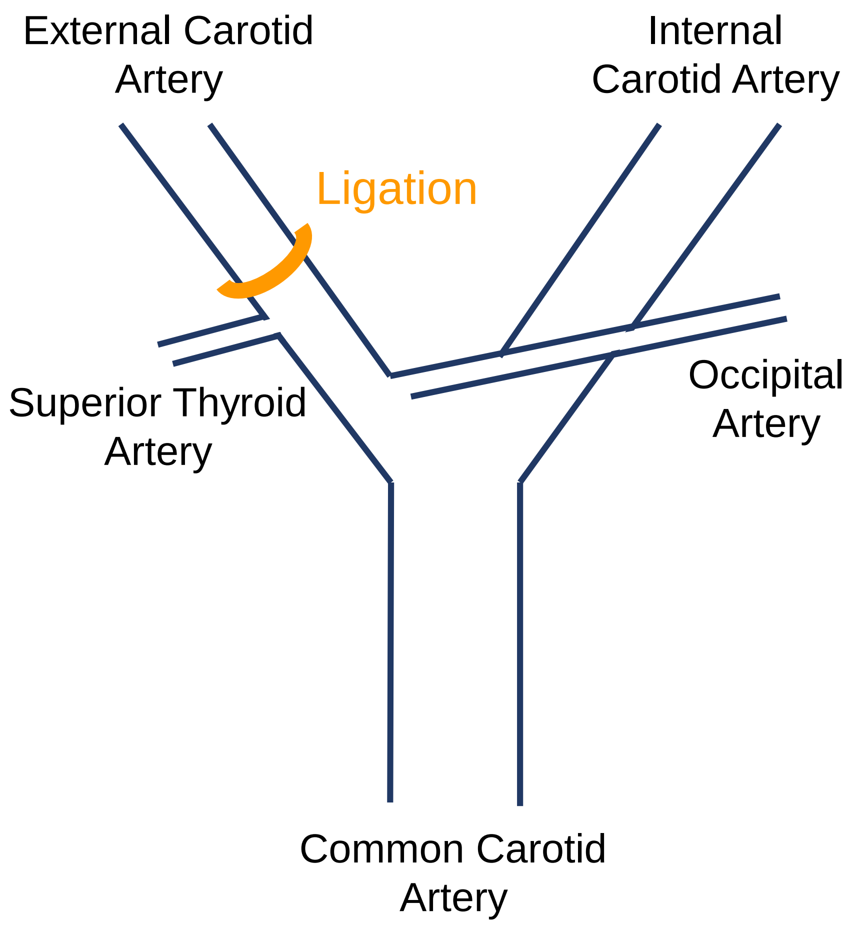


**Supplementary Figure 8. Schematic illustrating the site of partial ligation in the common carotid artery.**
